# Comparative larval single-cell transcriptomics provides insights into the evolutionary origins of sensory-neural programmes in Porifera and Cnidaria

**DOI:** 10.64898/2026.11.08.765225

**Authors:** Imran Luqman, Elizabeth Williams, Jordi Solana, Vengamanaidu Modepalli

## Abstract

Understanding how specialised cell types emerged and diversified is fundamental to reconstructing animal evolution. The origins of neural and sensory-secretory cell types are particularly intriguing, and early diverging non-neuronal metazoans, such as sponges, provide critical insights into pre-neural molecular toolkits. Free-swimming larval stages of sponges and cnidarians provide an unique context for investigating these questions, as these larvae rely on specialised sensory and signalling systems to detect environmental cues and coordinate behaviour during dispersal and settlement. Here, we integrated larval single-cell transcriptomic datasets from the demosponge *Amphimedon queenslandica* and four cnidarian species (*Nematostella vectensis*, *Astroides calycularis*, *Pocillopora acuta* and *Clytia hemisphaerica*). Cross-species mapping using Self-Assembling Manifold mapping (SAMap) recovered broad conservation of major cell-type identities among cnidarians, including neural, secretory, cnidocyte, epithelial, gastrodermal and apical populations. Extending these comparisons to *A. queenslandica* displayed transcriptional similarities between several sponge larval cell populations and cnidarian neural, sensory and progenitor-associated cell types. Within-species analyses showed that these cell populations remained transcriptionally integrated within the broader epithelial and sensory cell type of the sponge larva. Neural-associated gene modules, including conserved families associated with photoreception, neural fate specification and postsynaptic scaffolding, were distributed across distinct epithelial and sensory-like sponge larval cell populations rather than being restricted to a single proto-neural cell type. Our findings suggest that molecular components subsequently incorporated into neural systems were already deployed across diverse cell types in a common ancestor of sponges & cnidarians. Our work supports an evolutionary scenario in which extant neural and sensory-secretory cell types arose through the diversification and reorganisation of molecular programmes distributed across multiple ancestral sensory-secretory cell types.

## Introduction

Understanding how cell types emerged and diversified is central to reconstructing the evolution of animal complexity^1–3^. Early-diverging metazoans (Porifera, Cnidaria, Placozoa and Ctenophora) offer unique insights into this question, as they span critical evolutionary transitions associated with the emergence and diversification of key animal cell types^4–7^. These lineages exhibit markedly different levels of cellular and nervous-system organisation. For example, cnidarians and ctenophores possess diverse specialised neuronal cell types organised within a nervous system, whereas poriferans and placozoans lack neurons despite possessing specialised cell populations associated with sensory, secretory and signalling functions. Comparing cell-type organisation within and across these deeply diverged lineages therefore provides an opportunity to investigate the evolutionary relationships between non-neural sensory-secretory cells and specialised neuronal cell types.

While adult stages of early-diverging metazoans exhibit considerable diversity in body-plan and cellular organisation, the ciliated, free-swimming larval stages of cnidarians and poriferans (sponges) provide a broadly comparable developmental and ecological context in which specialised cell types must have adapted to similar environmental challenges. Whether the free-swimming larval stages of sponges and cnidarians are homologous, reflecting inheritance from a common ancestor, or evolved independently remains debated^8–10^. Nevertheless, these shared developmental and ecological demands provide an opportunity to investigate sensory-associated cell types and molecular programmes without presupposing homology between their larval stages or constituent cell types. Since successful habitat selection for larval settlement and metamorphosis is critical for subsequent survival and future reproductive success^11–13^, these larval stages are subject to strong selective pressures, particularly regarding sensory perception and coordinated regulation of behavioural outputs. Despite their relatively simple cellular organisation, both sponge and cnidarian larvae display sophisticated navigation behaviours in response to multiple environmental cues, such as light and chemicals, requiring the integration of sensory information with the modulation of specific cellular responses, such as ciliary beating^14–17^. These behavioural demands are supported by specialised sensory and signalling cell types that form a prominent component of sponge and cnidarian larval biology^16,18,19^.

Studies of cnidarian and sponge larvae have identified diverse sensory-secretory and neuronal cell types distributed throughout the larval epithelium^16,18,20^, often concentrated towards the anterior or apical region^21–24^, where they contribute to environmental sensing and the regulation of swimming and metamorphosis. Notably, some of these specialised sensory cell populations and structures are transient features of the larval stage and are lost or extensively remodelled during metamorphosis, including ciliated sensory and locomotory cells and sensory-secretory and neurosecretory cell populations^22,25–27^. Many, although not all, larval forms possess apical sensory organs containing ciliated sensory cells (Fig. 1a), together with peptidergic and secretory cell populations, which have been proposed to represent deeply conserved sensory structures across animal phyla^21,28–30^. Together, the diversity and specialised deployment of these larval sensory-associated cell types provide an informative context for investigating how sensory, secretory and neuronal cell identities have diversified during animal evolution.

**Figure 1.**
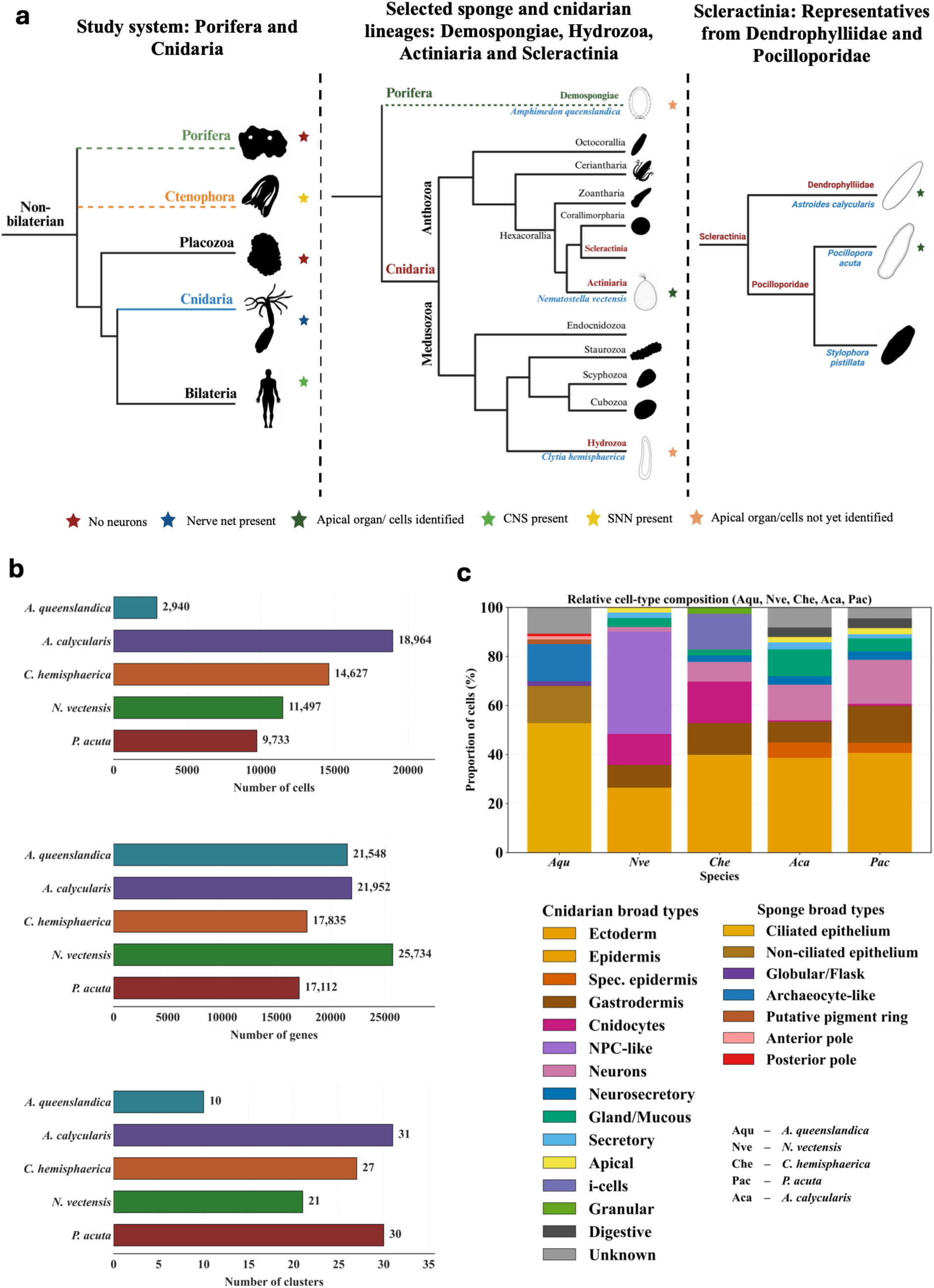
Study system and single-cell transcriptomic datasets analysed across five early-diverging metazoans. **(a)** Hierarchical phylogenetic overview of the species included in this study. From left to right, the figure highlights the placement of the sampled taxa within Metazoa, the diversification of major cnidarian lineages, and the relationships among the sampled scleractinian corals. The analyses include the demosponge *Amphimedon queenslandica* (Porifera), the hydrozoan *Clytia hemisphaerica*, the actiniarian *Nematostella vectensis*, and two scleractinian corals representing distinct families, *Astroides calycularis* (Dendrophylliidae) and *Pocillopora acuta* (Pocilloporidae). Coloured stars indicate nervous system and apical organ status: red, neurons absent; blue, neurons present (nerve net); yellow, possesses a syncytial nerve net (SNN); orange, no characterised apical organ/cells; dark green, characterised apical organ/cells; light green, possesses a central nervous system (CNS). **(b)** Summary of the five single-cell transcriptomic datasets showing the number of cells, detected genes and annotated cell clusters analysed for each species. **(c)** Relative proportions of cell assigned to harmonized broad cell type categories for each species, displayed as 100% stacked barplots. Cell-type proportions were calculated from the number of cells assigned to each category within the respective single-cell datasets.

Among sponges, the ciliated parenchymella larva of the demosponge *Amphimedon queenslandica* provides a unique system for cell type studies due to its history as a laboratory model for evolutionary and ecological developmental biology^31^ and is currently the only sponge species for which larval single-cell transcriptomic data are available^32^. *A. queenslandica* larvae respond to multiple environmental cues during their free-swimming phase, including light and biochemical settlement cues associated with suitable benthic substrates such as coralline algae^16,17,33^. These responses involve specialised cell populations distributed along the larval body^16^, including posterior pigment-ring cells associated with phototactic behaviour and anterior flask cells implicated in sensing settlement cues and initiating metamorphosis^17^, as well as vertically inherited bacterial symbionts that provide essential metabolic precursors for host nitric oxide signalling during settlement and metamorphosis^16,17,34^. Globular cells in this sponge larva have also been implicated in larval sensory and signalling functions, including nitric oxide signalling associated with settlement and metamorphosis^18,35^. The presence of postsynaptic scaffolding proteins and other molecular components associated with neural communication in sponges^36–38^ suggests that many of the molecular building blocks underlying neural communication were present prior to the evolution of neurons, with components of this machinery expressed in sponge larval sensory-secretory flask cells^16,36^.

In cnidarian larvae, many of the developmental and behavioural processes described above are regulated by neurons and neurosecretory cells, with neural and neuropeptidergic signalling contributing to ciliary locomotion, responses to environmental stimuli and settlement-associated behaviours ^20,36–40^. Among cnidarians, the sea anemone *Nematostella vectensis*, an anthozoan, has emerged as a major laboratory model for investigating neurogenesis and nervous-system development, with its planula larva possessing diverse sensory, neuronal and secretory cell populations^41–43^. These include a distinct apical sensory organ characterised by a prominent ciliary tuft^15,28^. Studies in *N. vectensis* have therefore provided detailed insight into the development and molecular organisation of the cnidarian larval nervous system^42,44,45^. In contrast, the hydrozoan *Clytia hemisphaerica*, another established laboratory model, lacks a transcriptionally distinct apical cell population equivalent to that identified in anthozoan larvae, instead possessing specialised sensory and neurosecretory cell populations organised across different regions of the aboral ectoderm^46^.

Despite growing evidence that many molecular components associated with neural communication predate canonical neurons^47–49^, the evolutionary relationships between non-neural sensory-secretory cell types and neuronal cell types remain poorly resolved, particularly how these cell types share conserved transcriptional programmes across early-diverging metazoans. Comparisons of cell-type transcriptional profiles between cnidarian and sponge larvae have the potential to provide unique insights into a central question of animal biology: the evolutionary history of neurons. Single-cell transcriptomic studies have increasingly characterised cellular diversity across early-diverging animal lineages, including sponges and cnidarians, generating cell-type atlases spanning both larval and adult developmental stages^32,41,50–52^. While many of these studies have focused on individual species, recent comparative analyses have begun to use single-cell transcriptomics to examine cell-type relationships across species. Comparative analysis of aboral cell types of larval *Clytia hemisphaerica* and corals (*Astroides calycularis* and *Pocillopora acuta*) identified conserved cell populations and signalling programmes across these three species, demonstrating the potential of cross-species single-cell approaches for resolving conserved features of larval cell-type organisation^46^. However, this study focused specifically on aboral cell populations within cnidarians and did not apply transcriptome-wide cross-species computationalmapping to systematically assess relationships across the full diversity of larval cell types or across more deeply diverged animal lineages.

Recent advances in single-cell transcriptomics^53–55^, combined with computational approaches such as Self-Assembling Manifold mapping (SAMap), permit systematic comparisons of cell-type transcriptional relationships and conserved molecular programmes across phylogenetically distant species^56^. The increasing availability of single-cell transcriptomic atlases across early-diverging metazoans now provides an opportunity to extend such comparisons across greater evolutionary distances and address a central question: how are sensory-secretory and neural-associated molecular programmes distributed across deeply diverged animal cell types? To address this question, we employed comparative single-cell transcriptomic analyses to investigate cell-type relationships across sponge and cnidarian larvae using previously published single-cell RNA-seq datasets from the demosponge *Amphimedon queenslandica* and four cnidarian species representing both anthozoan and hydrozoan lineages: *Nematostella vectensis*, *Clytia hemisphaerica*, *Pocillopora acuta* and *Astroides calycularis* (Fig. 1a). We integrated these larval single-cell atlases using cross-species cell-type mapping together with analyses of transcriptional relationships, transcriptional connectivity (inferred from graph-based relationships between cell populations) and gene co-expression programmes. These complementary approaches enabled us to identify shared molecular features across larval cell types and to investigate transcriptional relationships between sponge and cnidarian populations, with particular focus on sensory-secretory, neuropeptidergic and neural-associated cell types.

## Methods

### Single-cell transcriptomic datasets and preprocessing

Publicly available larval single-cell transcriptomic datasets from *Amphimedon queenslandica*^32^, *Nematostella vectensis*^41^, *Clytia hemisphaerica*, *Astroides calycularis* and *Pocillopora acuta*^46^ were analysed in Python using Scanpy (v1.11.5)^57^. Species-specific .h5ad objects containing previously quality-controlled and normalised single-cell transcriptomic data were used for all downstream analyses.

For *A. queenslandica*, the published larval dataset was re-clustered using the log-normalised expression matrix. The 3,000 most highly variable genes were identified and used for principal component analysis, neighbourhood graph construction and Leiden clustering across resolutions of 0.2–1.0. A resolution of 0.4 was selected as a balance between increased cellular resolution and preservation of the published metacell structure. Leiden clusters were annotated by comparison with the original published cell-type annotations using cluster-wise contingency tables. Two clusters associated with the original posterior-pole population were further distinguished using previously reported pigment-ring marker genes^49^, including *Ferritin* (Aqu2.1.42556_001 and Aqu2.1.42554_001), *PLOD* (Aqu2.1.27255_001), *Cry2* (Aqu2.1.32109_001), *NM Myhc* (Aqu2.1.43313_001), *ST Myhc* (Aqu2.1.43326_001) and *NK5/6/7b* (Aqu2.1.40608_001), scored using Scanpy’s score_genes function. The cluster showing stronger enrichment for this marker programme was annotated as a putative pigment-ring population.

For *N. vectensis*, Leiden clusters generated at a resolution of 0.6 were used for downstream analyses. The resulting clusters were annotated by comparison with the published cell-type classifications using cluster-wise contingency tables, with refined cell-type annotations assigned according to their correspondence with the original annotations. The apical-organ population was further identified using a previously reported 15-gene apical-organ marker set^15,28^, with marker enrichment calculated across clusters using Scanpy’s score_genes function and the highest-scoring cluster assigned as the apical organ. Published cluster annotations were otherwise retained for *C. hemisphaerica*, *A. calycularis* and *P. acuta*.

### Cell-state relationship and graph abstraction analyses

Species-specific dendrograms were generated using Scanpy (sc.tl.dendrogram) to examine transcriptional relationships among annotated cell populations in *A. queenslandica*, *N. vectensis*, *C. hemisphaerica*, *P. acuta* and *A. calycularis*. The existing Leiden cluster assignments were used throughout (leiden_0.6 for *N. vectensis*, leiden_new for *C. hemisphaerica* and *A. calycularis*, and leiden for *P.acuta*), together with the custom leiden_r0.4 clustering generated for *A. queenslandica*. Dendrograms were computed from the principal component representations contained within each dataset. For visualisation, clusters were coloured according to manually assigned broad cell-type annotations.

Partition-based Graph Abstraction (PAGA)^58^ was performed using the corresponding species-specific cluster annotations to infer large-scale transcriptional connectivity among annotated cell populations. *A. queenslandica* was analysed using the custom leiden_r0.4 clustering, with PAGA connectivity projected onto the published larval two-dimensional coordinate system and additionally visualised using force-directed spring layouts. PAGA graphs were set with a threshold to retain higher-confidence relationships and coloured according to manually assigned broad cell-type annotations.

### Cross-species cell-state mapping using SAMap

Cross-species gene homology maps were generated using the BLAST-based mapping workflow implemented in the original SAMap pipeline^56^. Species-specific protein databases were constructed where required, and reciprocal BLASTP searches were performed between all species pairs to identify putative homologous genes for downstream comparative analyses. BLAST searches were conducted using a maximum of one high-scoring segment pair per query (max_hsps = 1) and an e-value threshold of 1 × 10⁻⁶. Pairwise alignments were exported in tabular format (outfmt 6) and used to generate bidirectional gene homology maps for all species pairs prior to SAMap integration.

Cross-species cell-state relationships were inferred using SAMap following the original published workflow^56^. Single-cell transcriptomic datasets from *A. queenslandica*, *N. vectensis*, *C. hemisphaerica*, *P. acuta* and *A. calycularis* were integrated using pairwise manifold alignment (pairwise = True) to identify transcriptionally similar cell populations across species. The resulting integrated SAMap object was retained for all downstream comparative analyses.

To preserve biologically curated cell-state identities, the standard downstream SAMap mapping workflow was modified to retain the original species-specific cluster annotations rather than SAMap-derived reclustering. Mapping analyses therefore used the manually curated cluster identities corresponding to leiden_new for *A. calycularis* and *C. hemisphaerica*, leiden for *P. acuta*, leiden_0.6 for *N. vectensis*, and the custom leiden_r0.4 annotation generated for *A. queenslandica*. Cross-species mapping scores were extracted using the get_mapping_scores function from samap.analysis, retaining the complete set of pairwise cluster relationships (n_top = 0) to generate transcriptional similarity matrices for downstream comparative analyses and network visualisation.

### Multi-species radial cell-type tree construction

A multi-species cell-type tree was constructed to visualise cross-species transcriptional relationships among annotated cell populations from *Amphimedon queenslandica*, *Nematostella vectensis*, *Clytia hemisphaerica*, *Pocillopora acuta* and *Astroides calycularis*. The SAMap cluster similarity matrix was used as the input, with each annotated cluster represented by its transcriptional similarity profile across all other annotated cell populations. All annotated clusters, including the custom *A. queenslandica* leiden_r0.4 clusters, were included in the analysis.

Pairwise similarities between cluster similarity profiles were converted to correlation distances and subjected to hierarchical clustering using the unweighted pair-group method with arithmetic mean (UPGMA; average linkage)^59^. The resulting hierarchical tree was rendered as a radial dendrogram using custom Python scripts incorporating pandas, NumPy, SciPy and Matplotlib.

### Graph-based SAMap network visualisation

Graph-based representations of cross-species cell-state relationships were constructed from the SAMap cluster similarity matrices using custom Python workflows. Annotated cell clusters were represented as nodes and pairwise SAMap mapping scores as weighted interspecies edges, with within-species connections excluded. To retain the strongest transcriptional relationships, top-*k* filtering was applied to preserve the highest-scoring cross-species connection for each cluster– species pair. Networks were generated using *k* = 2 for the cnidarian-only analysis and *k* = 1 for the cnidarian–*Amphimedon queenslandica* analysis. Similarity thresholds of 0.075, 0.10 and 0.15 were evaluated for the cnidarian-only network, while thresholds of 0.10, 0.15, 0.20, 0.25 and 0.30 were evaluated for the cnidarian–*A. queenslandica* network. Unless otherwise stated, the primary analyses used thresholds of 0.075 and 0.15, respectively. Equivalent networks generated across all tested thresholds are provided in the Supplementary Information. The custom *A. queenslandica* leiden_r0.4 clusters were used throughout analyses incorporating sponge cell populations.

Network communities were identified using greedy modularity optimisation and visualised using force-directed layouts. Community-level summary networks were generated using the igraph cluster graph framework, in which communities were contracted into single meta-nodes.

### Reciprocal best-hit BLAST analyses

Putative orthologues of selected genes were identified using reciprocal best-hit BLASTP analyses of their encoded proteins. Protein sequences corresponding to these query genes were searched against the corresponding predicted proteomes using BLASTP (NCBI BLAST+ v2.16.0), and the highest-scoring hit was retained (e-value threshold = 1 × 10⁻⁵). The corresponding protein sequence was then extracted and searched reciprocally against a BLAST database constructed from the original query protein sequence(s). Genes were inferred to be reciprocal best-hit orthologues only when the highest-scoring hit in the reciprocal search matched the original query protein sequence.

### Identification of putative neural progenitor cell (NPC)-like populations

Putative neural progenitor cell (NPC)-like populations in cnidarians, and sponge cell populations expressing homologues of cnidarian NPC markers, were identified across the single-cell transcriptomic datasets using a comparative marker-based approach. Candidate NPC marker genes were obtained from molecularly characterised NPC populations in *Nematostella vectensis*^41^. The NPC-associated marker genes used for downstream analyses were: PCNA-like (NVE26120); NvNeurogenin1 (NVE6000); NvSoxC (NVE12762); DLGAP5-like (NVE7848); SMC4-like (NVE24422); NvMyc3 (NVE20961); RIR1-like (NVE8178); SMC2-like (NVE8734); RIR2-like (NVE21390); NvMyc1 (NVE20962); CENPE-like (NVE16653); NvSoxB.2 (NVE24655); CENPF-like (NVE9959); and NvNanos1 (NVE13963). Species-specific NPC marker gene sets were compiled by identifying annotated homologues in *C. hemisphaerica*, *P. acuta*, *A. calycularis* and *A. queenslandica*. Where annotated homologues were unavailable, reciprocal best-hit BLASTP analyses were performed using the previously described BLAST parameters. Annotation-based matches and reciprocal best-hit orthologues were combined to generate species-specific NPC marker gene sets. NPC-associated gene expression was examined using Scanpy dot plots, and module scores were calculated for each cell using sc.tl.score_genes. Mean cluster-level module scores were used to identify candidate NPC-like populations within each species.

Candidate NPC-like populations were compared across species using SAMap-derived mapping scores, with the previously characterised *N. vectensis* NPC populations serving as the reference. Pairwise mapping matrices were queried to identify the strongest cross-species transcriptional similarities.

### Identification of putative sensory-neural populations in *Amphimedon queenslandica*

To identify *A. queenslandica* cell populations expressing ancestral sensory-neural signatures, the neural Gene Ontology (GO), synaptic, signalling pathway and homeobox transcription factor gene sets were compiled from previously published studies describing sponge sensory biology, neural-like cell types and conserved neural-associated gene repertoires, with particular emphasis on studies of *Amphimedon*. Gene module enrichment was assessed across the custom leiden_r0.4 cell clusters using Scanpy. Module scores were calculated for individual cells using sc.tl.score_genes, and cluster-level enrichment was quantified using the mean module score for each cluster. The highest-scoring clusters for each gene module were visualised on UMAP embeddings.

To evaluate whether module enrichments were greater than expected by chance, module-score distributions within each cluster were first assessed for normality using the Shapiro–Wilk test. As most module-score distributions deviated from normality, non-parametric statistical analyses were performed. For each gene module, module scores within each cluster were compared against all remaining cells using one-sided Mann–Whitney U tests, with *P*-values corrected for multiple testing using the Benjamini–Hochberg false discovery rate (FDR) procedure. Statistical significance was further assessed using a permutation-based null model, in which cluster labels were randomly permuted 1,000 times while preserving cluster size. For each permutation, the mean module score of the focal cluster was recalculated to generate a null distribution, from which empirical *P*-values and permutation *Z*-scores were derived.

Clusters showing statistically supported enrichment for one or more neural-associated gene modules were subsequently compared with the cross-species SAMap cluster similarity network. The highest-ranking *Amphimedon* clusters were examined for their transcriptional relationships with cnidarian cell populations, with particular emphasis on neural, neurosecretory, gland-mucous, secretory, cnidocyte and apical organ cell types, to determine whether sponge populations enriched for neural-associated gene programmes also exhibited conserved cross-species transcriptional similarity.

### Weighted Gene Co-expression Network Analysis (WGCNA)

Weighted Gene Co-expression Network Analysis (WGCNA) was performed independently for *A. queenslandica*, *N. vectensis*, *C. hemisphaerica*, *P. acuta* and *A. calycularis* using the PyWGCNA framework^60^. Pseudobulk expression matrices were generated by averaging log-normalised gene expression across sample–cluster combinations, and genes expressed in fewer than 10% of pseudobulk samples were excluded. Co-expression networks were constructed using PyWGCNA, including soft-threshold selection, module detection, and calculation of module eigengenes and module membership values (kME). The top 50 hub genes from each module were retained for downstream analyses, excluding the unclassified grey module. Modules were manually annotated according to the predominant functions of their hub genes using UniProt annotations and visualised as row-scaled module eigengene heatmaps across annotated cell populations.

To compare co-expression programmes across species, homologous gene relationships were obtained from the SAMap BLAST-derived orthology mappings and clustered into species-independent orthogroups using a union–find algorithm. Species-specific WGCNA modules were mapped to these orthogroups, and pairwise module similarity was quantified using the Jaccard index, with statistical significance assessed using the hypergeometric test. Conserved module associations were retained using thresholds of Jaccard similarity ≥ 0.05 and hypergeometric *P* < 0.01, and visualised as weighted module networks and heatmaps of Jaccard similarity and −log10-transformed hypergeometric *P*-values.

### Data availability

All custom scripts used for data preprocessing, cross-species transcriptomic analyses and figure generation are available through the associated GitHub repository [https://github.com/ImLuq14/Sponge-cnidarian-cross-species-comparisons]. Processed single-cell datasets (.h5ad files), FASTA files used in the analyses and supporting data are available on Zenodo [https://doi.org/10.5281/zenodo.23942514]. Additional methodological details and supporting analyses are provided in the Supplementary Methods, Figures and Tables where indicated.

## Results

### Integration of larval single-cell transcriptomic datasets to reveal larval cell-types across Porifera and Cnidaria

To investigate the evolutionary relationships between larval sponge and cnidarian cell types, we re-analysed published single-cell transcriptomic atlases from the demosponge *Amphimedon queenslandica* and four cnidarian species: *Nematostella vectensis*, *Clytia hemisphaerica*, *Astroides calycularis*, and *Pocillopora acuta* (Fig. 1a; Supplementary Fig. S1). Processed Scanpy objects were obtained for the cnidarian species, whereas the *A. queenslandica* dataset was re-analysed from the published expression data. These datasets provided the basis for subsequent cross-species comparative analyses (Supplementary Fig. S1).

The five atlases varied substantially in size and complexity (Fig. 1b). Cell numbers ranged from 2,940 in *A. queenslandica* to 18,964 in *A. calycularis*, while detected genes ranged from 17,112 in *P. acuta* to 25,734 in *N. vectensis*. Cluster numbers ranged from 10 in *A. queenslandica* to 31 in *A. calycularis*, with the cnidarian datasets containing 21–31 annotated clusters, reflecting greater cellular complexity than the sponge atlas. To facilitate cross-species comparisons, clusters were grouped into broad cell-type categories based on their original annotations (Fig. 1c). Where populations retain a “neurosecretory” annotation from the source datasets, this term is used as an original cell-type label and should be interpreted broadly as referring to secretory populations rather than as evidence of demonstrated neurosecretory function. In cnidarians, epidermal, gastrodermal, neuronal, and cnidocyte populations accounted for substantial proportions of the sampled cells, with neuronal diversity ranging from five clusters in *C. hemisphaerica* to nine clusters in *P. acuta*. Neurosecretory, gland/mucous, secretory, and apical populations were also identified across cnidarians. The *A. queenslandica* atlas comprised ciliated and non-ciliated epithelial cells, archaeocyte-like cells, previously described sensory-like globular/flask and pigment ring cells, plus anterior and posterior pole cells.

Marker-based analyses further supported the refined annotation of specific sensory-associated populations in *N. vectensis* and *A. queenslandica* (Supplementary Fig. S2). In *N. vectensis*, the apical-organ marker programme was most strongly enriched in cluster 14, supporting its annotation as the apical-organ population. In *A. queenslandica*, pigment-ring markers showed stronger enrichment in cluster 4 than in cluster 8, the two clusters corresponding to the original posterior-pole population, supporting the annotation of cluster 4 as a putative pigment-ring population. Together, these analyses established a consistently annotated set of larval cell populations across the five species, allowing subsequent cross-species comparisons of cell-type transcriptional relationships.

### Dendrogram and PAGA analyses resolve transcriptional relationships among larval cell types

To identify shared cell types across cnidarian larvae, we combined dendrogram and PAGA analyses, which recovered highly similar transcriptional organisation across the four cnidarian larval atlases (Fig. 2a–h). In all species, neuronal and secretory populations showed close transcriptional relationships rather than forming completely separate branches. PAGA supported these relationships, revealing extensive connectivity among neuronal and secretory populations, with central nodes corresponding to neuronal populations in *A. calycularis*, *P. acuta* and *C. hemisphaerica*, and to neurogenic progenitors in *N. vectensis*. Gland/mucous and other secretory populations generally formed neighbouring but transcriptionally distinct modules that retained connections to the broader neural-associated network, whereas apical cell populations were consistently associated with epidermal cell types. *N. vectensis* additionally resolved a progenitor-centred organisation linking neurogenic progenitors, secretory populations, sensory-associated apical cells and neurons, while *C. hemisphaerica* contained a distinct granular secretory lineage separate from the neural-associated network.

**Figure 2.**
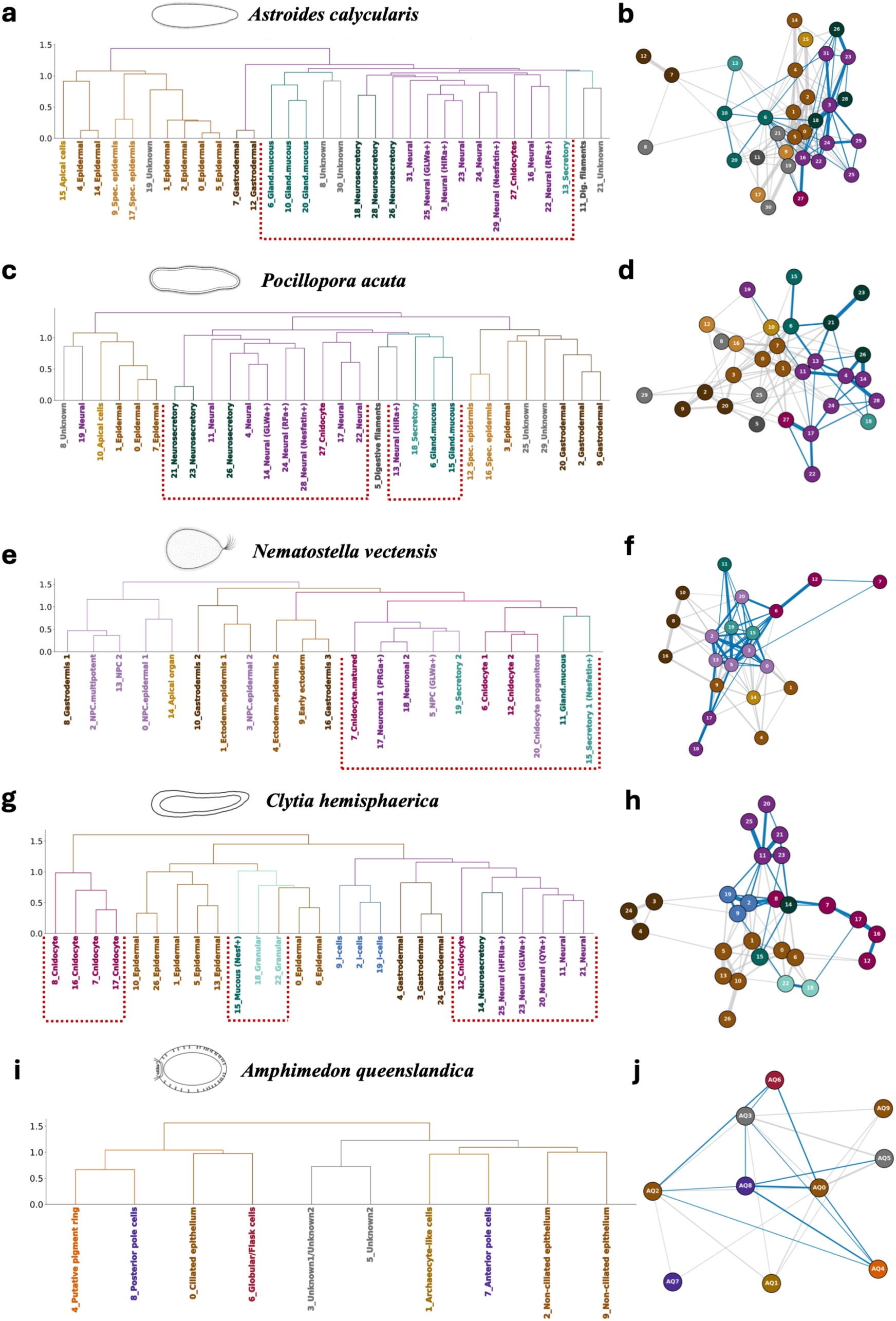
Within-species hierarchical dendrograms and partition-based graph abstraction (PAGA) networks. (a, c, e, g, **i)** Hierarchical dendrograms constructed from pseudobulk transcriptomic profiles, illustrating transcriptomic similarity among annotated cell clusters in **(a)** *A. calycularis*, **(c)** *P. acuta*, **(e)** *N. vectensis*, **(g)** *C. hemisphaerica* and **(i)** *A. queenslandica*. Dendrogram branches and cluster labels are coloured according to harmonised broad cell-type annotations. Red dotted boxes in highlight groups of neuronal and sensory-secretory cell populations exhibiting close transcriptional relationships. **(b, d, f, h, j)** Corresponding partition-based graph abstraction (PAGA) networks inferred from the same datasets. Nodes represent annotated cell clusters and are coloured according to the same broad cell-type annotations. Edges indicate inferred transcriptional connectivity between clusters, with thicker and darker edges representing stronger connectivity. Blue edges denote connections between neural cell clusters and other sensory- and secretory-associated cell clusters and are shown for emphasis.

In the case of the *A. queenslandica* atlas, hierarchical clustering identified a close relationship between pigment ring cells (cluster 4) and posterior pole cells (cluster 8), which subsequently grouped with ciliated epithelial cells (cluster 0) and globular/flask cells (cluster 6), forming a broader epithelial-associated assemblage. Globular/flask cells were most closely associated with ciliated epithelial cells, whereas non-ciliated epithelial clusters 2 and 9 grouped with anterior pole cells (cluster 7) and archaeocyte-like cells (cluster 1). The putative sensory-secretory larval cell populations were placed within a broader epithelial transcriptional landscape (Fig. 2i, j). PAGA supported this organisation, revealing direct connectivity between putative sensory-secretory and epithelial populations. Posterior pole, pigment ring and globular/flask cells are all connected to epithelial populations, with globular/flask cells linking both ciliated and non-ciliated epithelial states. Overall, these analyses revealed distinct patters of transcriptional organization, with cnidarian neural-associated populations forming closely connected groups, while sponge sensory-associated populations remained closely associated with epithelial cell types.

### BLAST-derived gene homology networks establish a framework for cross-species transcriptomic mapping

To assess the shared gene homology underlying the cross-species comparisons, we examined the distribution of BLAST-connected gene groups across the five species used for SAMap analysis (Fig. 3a–c). A total of 1,816 BLAST-connected gene groups were shared across all five species, while an additional 482 gene groups were shared exclusively among the four cnidarians. Among the cnidarians, 908 gene groups were shared exclusively among the three anthozoans (*A. calycularis*, *P. acuta* and *N. vectensis*), while the sister species *P. acuta* and *A. calycularis* exhibited the largest exclusive pairwise intersection, comprising 374 gene groups (Fig. 3a). Pairwise Jaccard similarity was highest among the three anthozoans, particularly between *P. acuta* and *A. calycularis* (0.89), followed by *N. vectensis* and *P. acuta* (0.85) and *N. vectensis* and *A. calycularis* (0.81). *C. hemisphaerica* showed lower similarity to the anthozoans (0.61–0.66), while *A. queenslandica* showed its highest similarity with *C. hemisphaerica* (0.76) and lower similarity with the three anthozoans (0.48–0.52) (Fig. 3b). Between 13,792 and 21,327 genes per species were incorporated into the BLAST-derived homology graph used for SAMap, corresponding to 69.0–98.2% of the genes represented in the respective larval single-cell datasets (Fig. 3c). These results indicate that BLAST-derived gene homology patterns broadly reflect the phylogenetic relationships among the sampled species, while identifying a shared set of genes suitable for subsequent cross-species transcriptomic mapping.

**Figure 3.**
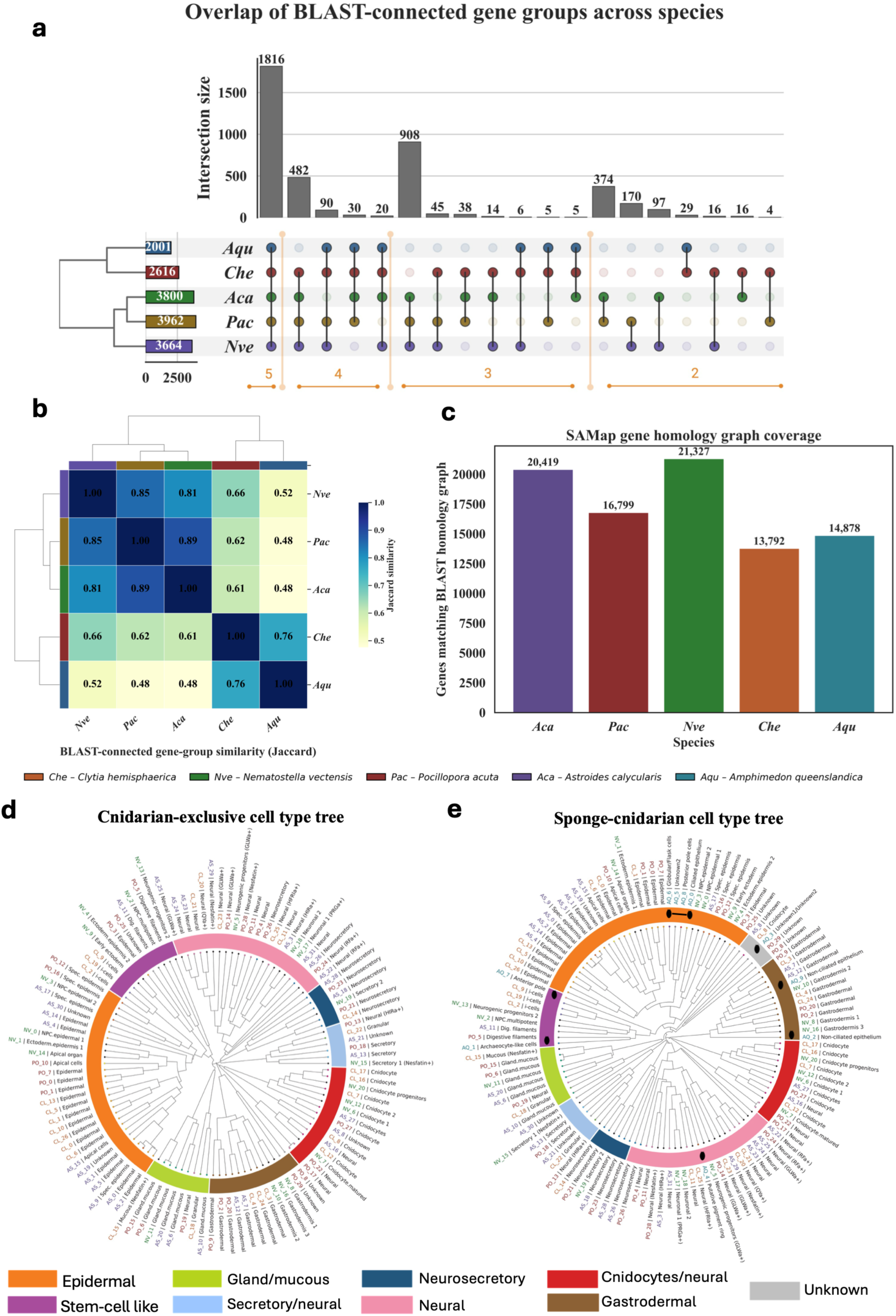
Overview of cross-species gene homology and cell-type relationships used for comparative analyses. **(a)** UpSet plot summarising the overlap of BLAST-connected gene groups across *A. calycularis*, *P. acuta*, *N. vectensis*, *C. hemisphaerica* and *A. queenslandica*. Horizontal bars indicate the total number of BLAST-connected gene groups identified in each species, while vertical bars show the size of each intersection across species combinations. **(b)** Pairwise Jaccard similarity heatmap of BLAST-connected gene groups, illustrating the extent of shared homologous gene content between species. **(c)** Number of genes from each species incorporated into the SAMap BLAST homology graph and used for cross-species mapping. **(d)** Radial hierarchical dendrogram of annotated cell clusters from the four cnidarian species based on SAMap-derived cell-type similarity profiles, coloured according to harmonised broad cell-type annotations. **(e)** Radial hierarchical dendrogram of annotated cell clusters from the four cnidarian species together with *A. queenslandica*, based on SAMap-derived cell-type similarity profiles and coloured according to harmonised broad cell-type annotations. Cluster labels are prefixed with species identifiers, and black larval silhouettes indicate the positions of *A. queenslandica* cell clusters.

### SAMap identifies conserved cross-species relationships among sponge and cnidarian larval cell types

Hierarchical clustering of SAMap-derived cell-type similarity profiles across the four cnidarian species resolved cell populations primarily according to broad cell identity rather than species origin (Fig. 3d). Neural populations formed a prominent cross-species clade comprising neuronal cell types from *A. calycularis*, *C. hemisphaerica*, *N. vectensis* and *P. acuta*, while neurosecretory populations formed a closely related but distinct assemblage. Additional clades broadly separated epithelial and apical populations, gastrodermal cells, gland/mucous-associated populations and cnidocytes, with representatives from multiple species recovered within each group. Together, these patterns demonstrate extensive conservation of broad transcriptional cell-type identities across the sampled cnidarians despite their phylogenetic divergence.

To investigate transcriptomic relationships among cell types across species, hierarchical clustering of SAMap-derived cell-type similarity profiles was visualised as a radial cell-type tree (Fig. 3e). Within this framework, *Amphimedon queenslandica* putative pigment ring cells (AQ4) clustered within the neural-associated clade, whereas globular/flask (AQ6), posterior pole (AQ8), and anterior pole (AQ7) cells grouped within the epithelial/apical assemblage (Fig. 3e).

To further characterise these relationships, we constructed a SAMap cluster similarity network comprising 110 transcriptionally defined clusters from the four cnidarian species (Fig. 4a). Network topology was robust across similarity thresholds (Supplementary Fig. S3), and a threshold of 0.075 was selected for subsequent analyses. Clusters grouped into ten conserved communities corresponding primarily to major cell-type identities rather than species, including epidermal, neural, gastrodermal, cnidocyte, gland/mucous, secretory, and neurosecretory populations. To examine transcriptional relationships among the broader cell-type communities, we contracted the network by representing each community as a single node (Fig. 4b). Additional comparisons across similarity thresholds and top-k partner settings showed broadly similar community-level organisation (Supplementary Fig. S4). The principal neural community occupied a central position and connected directly to secretory, neurosecretory, gland/mucous, and i-cell-associated communities, whereas epidermal and gastrodermal communities remained comparatively distinct. A smaller neural community formed an additional bridge to the cnidocyte community.

**Figure 4.**
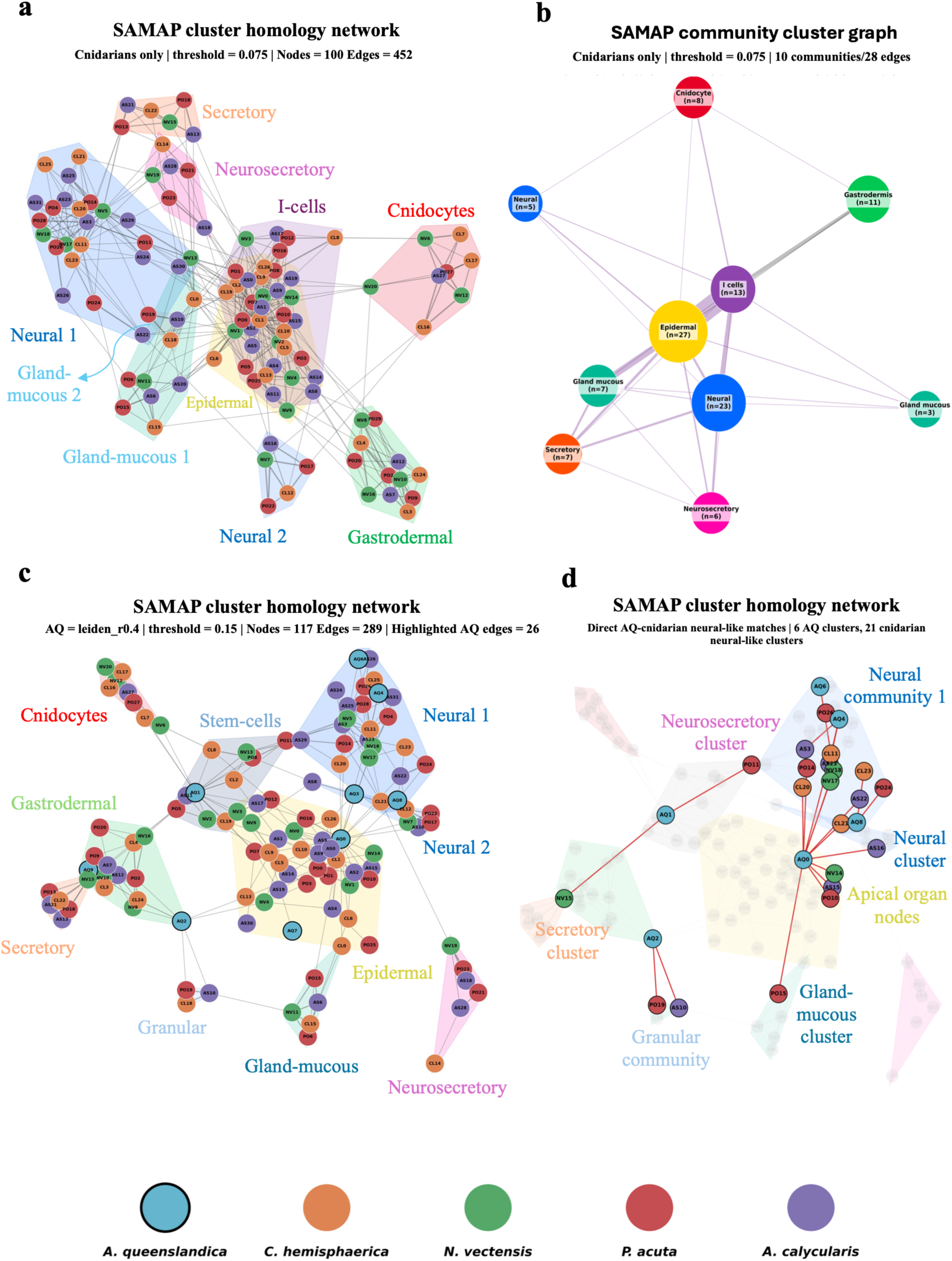
Cross-species SAMap cluster homology networks identify conserved cell-type communities and sponge–cnidarian relationships. **(a)** SAMap cluster homology network constructed from the four cnidarian species (*A. calycularis*, *P. acuta*, *N. vectensis* and *C. hemisphaerica*). Nodes represent annotated cell clusters, with node colours indicating species, and edges represent transcriptional similarity inferred by SAMap. The network was filtered using a similarity threshold of 0.075 while retaining the two strongest interspecies connections per cluster. Shaded polygons outline communities identified by greedy modularity analysis and are labelled according to their predominant broad cell-type identity; polygon colours distinguish the different cell-type communities. **(b)** Contracted community graph generated from the network shown in (a) using the igraph cluster graph algorithm. Nodes represent communities, with node colours corresponding to the cell-type communities shown in (a), node size proportional to the number of constituent clusters and edge width proportional to the cumulative similarity between connected communities. **(c)** SAMap cluster homology network including *A. queenslandica*. The network was filtered using a similarity threshold of 0.15 while retaining the strongest interspecies connection per cluster. Node and polygon colours follow the same conventions as in (a), with *A. queenslandica* represented by an additional node colour. **(d)** Simplified view of the network shown in (c), with unrelated connections greyed out to emphasise direct homology relationships between *Amphimedon* cell clusters and cnidarian neural, neurosecretory, gland-mucous and secretory communities.

To determine how sponge larval cell populations relate to these conserved cnidarian communities, *A. queenslandica* clusters were incorporated into the SAMap network (Fig. 4c). A similarity threshold of 0.15 retained a single connected network containing nine of the ten annotated sponge clusters (Supplementary Fig. S5; Supplementary Table 1). Rather than forming a sponge-specific community, sponge cell populations were distributed among conserved cnidarian communities according to transcriptional similarity. Ciliated epithelium and anterior pole cells grouped with the epidermal community, non-ciliated epithelial populations grouped with the gastrodermal community, and archaeocyte-like cells associated with a community containing i-cells, digestive filament populations, and neurogenic progenitors.

To examine the cross-species relationships underlying these community assignments, we visualised only the connections between *A. queenslandica* clusters and cnidarian neural-, neurosecretory-, secretory-, gland/mucous-, and apical-associated populations (Fig. 4d). Putative pigment ring, globular/flask, and posterior pole cells connected predominantly to the principal neural community, whereas ciliated epithelial cells formed multiple direct links to neural, neurosecretory, gland/mucous, and apical populations despite remaining within the epidermal community. Archaeocyte-like and non-ciliated epithelial populations exhibited comparatively fewer highlighted connections. These findings indicate that sponge larval cell populations exhibit transcriptional similarities to multiple cnidarian cell-type groups, including neural-associated populations, rather than forming a distinct sponge-specific group in the comparative network.

### Conserved progenitor-like cell states connect cnidarian neurogenic populations and sponge archaeocytes

To identify neurogenic progenitor cell (NPC)-like populations across species, genes enriched in the established *Nematostella vectensis* neurogenic progenitor cluster (NV13) were used to generate species-specific NPC scores based on reciprocal BLAST homologs (Fig. 5a). NPC-associated genes, including regulators of proliferation and developmental patterning such as *pcna, myc, nanos, sox* family genes, *dlgap5, cenpe/cenpf,* and *rnr1*, were enriched within restricted cell populations in each species. Ranking clusters by composite NPC score identified the highest-scoring candidate NPC-like population in each dataset: specialised epidermis (AS17) in *Astroides calycularis*, neurosecretory cells (PO11) in *Pocillopora acuta*, i-cells (CL9) in *Clytia hemisphaerica*, and archaeocyte-like cells (AQ1) in *Amphimedon queenslandica*.

**Figure 5.**
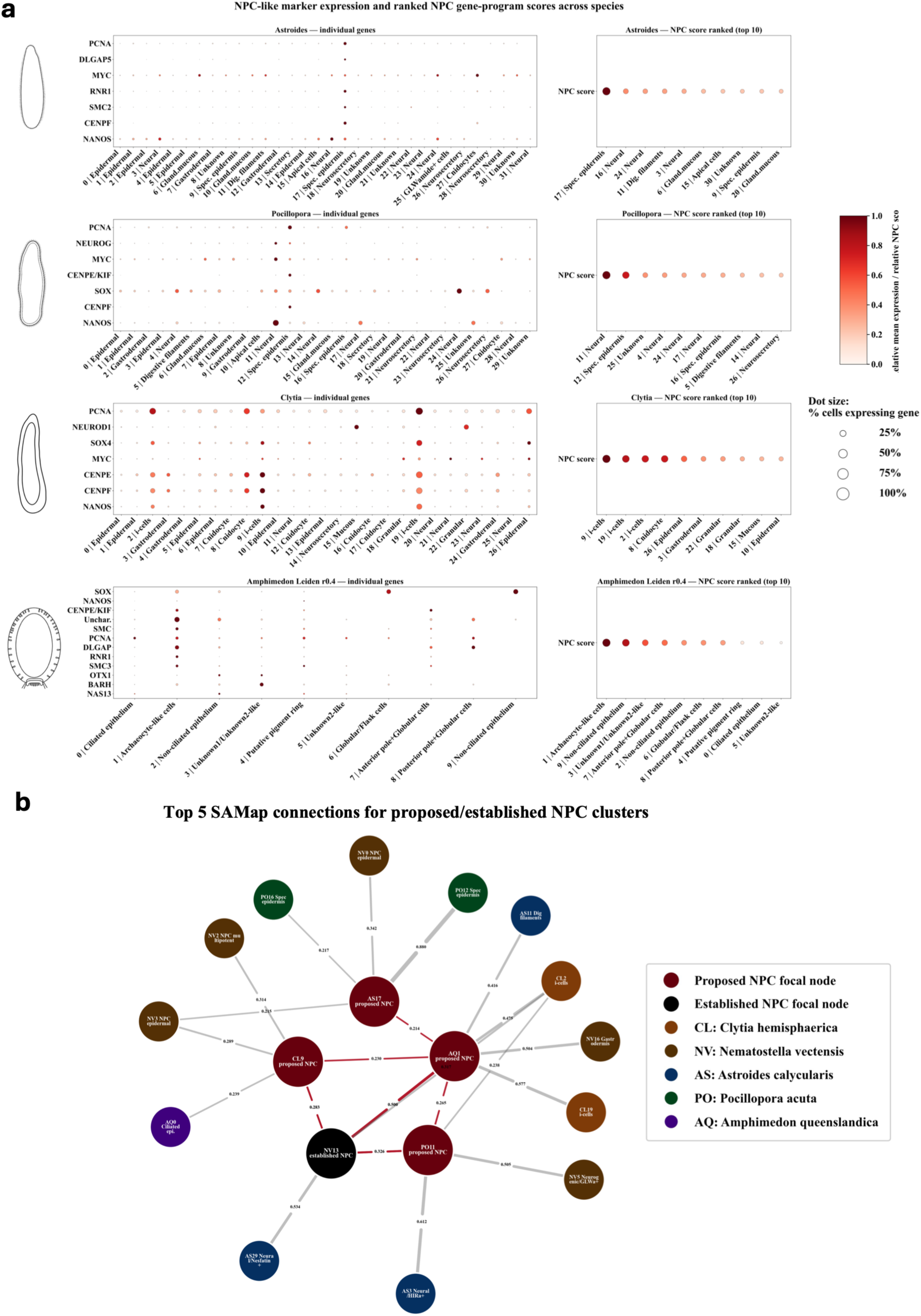
Identification of putative neural progenitor cell (NPC) populations across species using conserved *Nematostella* NPC markers. **(a)** Expression of conserved *Nematostella vectensis* NPC marker genes across annotated cell clusters in *A. calycularis*, *P. acuta*, *C. hemisphaerica* and *A. queenslandica*. For each species, the left dot plot shows the expression of individual NPC marker genes across cell clusters, where dot size represents the percentage of expressing cells and colour indicates scaled mean expression. The right dot plot shows the corresponding aggregate NPC gene-program score for each cluster, with clusters ranked from highest to lowest score to identify putative NPC populations. **(b)** SAMap network showing the five strongest cross-species homology relationships for proposed NPC populations identified in (a). Red nodes indicate proposed NPC clusters identified by marker-gene scoring, while the black node denotes the established *Nematostella* NPC population used as the reference. Edges represent SAMap similarity scores between proposed and established NPC populations and their closest homologous cell clusters across species.

To examine the relationships among these candidate NPC populations, the highest-scoring clusters were compared using SAMap (Fig. 5b). The established *Nematostella* neurogenic progenitor cluster (NV13) formed direct cross-species connections with each candidate NPC population, including AQ1 (SAMap score = 0.500), PO11 (0.326), CL9 (0.283), and AS17 (0.066). The archaeocyte-like cluster (AQ1) also connected directly to all proposed cnidarian NPC populations, while CL9 and AS17 showed additional links to progenitor- and stem-cell-associated populations, including NPC transition states, multipotent progenitors, and i-cells. Together, these findings suggest that progenitor-like cell populations in cnidarian and sponge larvae share transcriptional similarities, potentially reflecting common cellular programmes associated with proliferation and developmental plasticity.

### Independent neural gene sets identify sensory-like sponge cell populations linked to cnidarian neural cell types

To assess whether any *Amphimedon queenslandica* larval cell populations exhibit enrichment of neural-associated transcriptional programmes, we compiled gene modules representing sensory/neural, signalling pathway, synaptic, homeobox transcription factor, and neural Gene Ontology (GO) functions from three studies that identified candidate neural-like gene repertoires in sponges^49,61,62^ (Fig. 6a). Module-score distributions were first assessed for normality using the Shapiro–Wilk test and were found to be predominantly non-normal, supporting the use of non-parametric statistical analyses. Mann–Whitney U tests and permutation-based null models, with multiple-testing correction applied to both analyses, identified significant enrichment of sensory/neural genes in ciliated epithelium (cluster 0), non-ciliated epithelium (cluster 2) and Unknown1/Unknown2 (cluster 3); neural GO terms and synaptic genes in non-ciliated epithelium (cluster 2); homeobox transcription factors in ciliated epithelium (cluster 0), Unknown1/Unknown2 (cluster 3) and globular/flask cells (cluster 6); and signalling pathway genes in ciliated epithelium (cluster 0), non-ciliated epithelium (cluster 2), Unknown1/Unknown2 (cluster 3), putative pigment ring cells (cluster 4) and globular/flask cells (cluster 6). These findings demonstrate that neural-, sensory-, synaptic-, developmental- and signalling-associated transcriptional programmes are distributed across multiple larval cell populations rather than being restricted to a single cluster.

**Figure 6.**
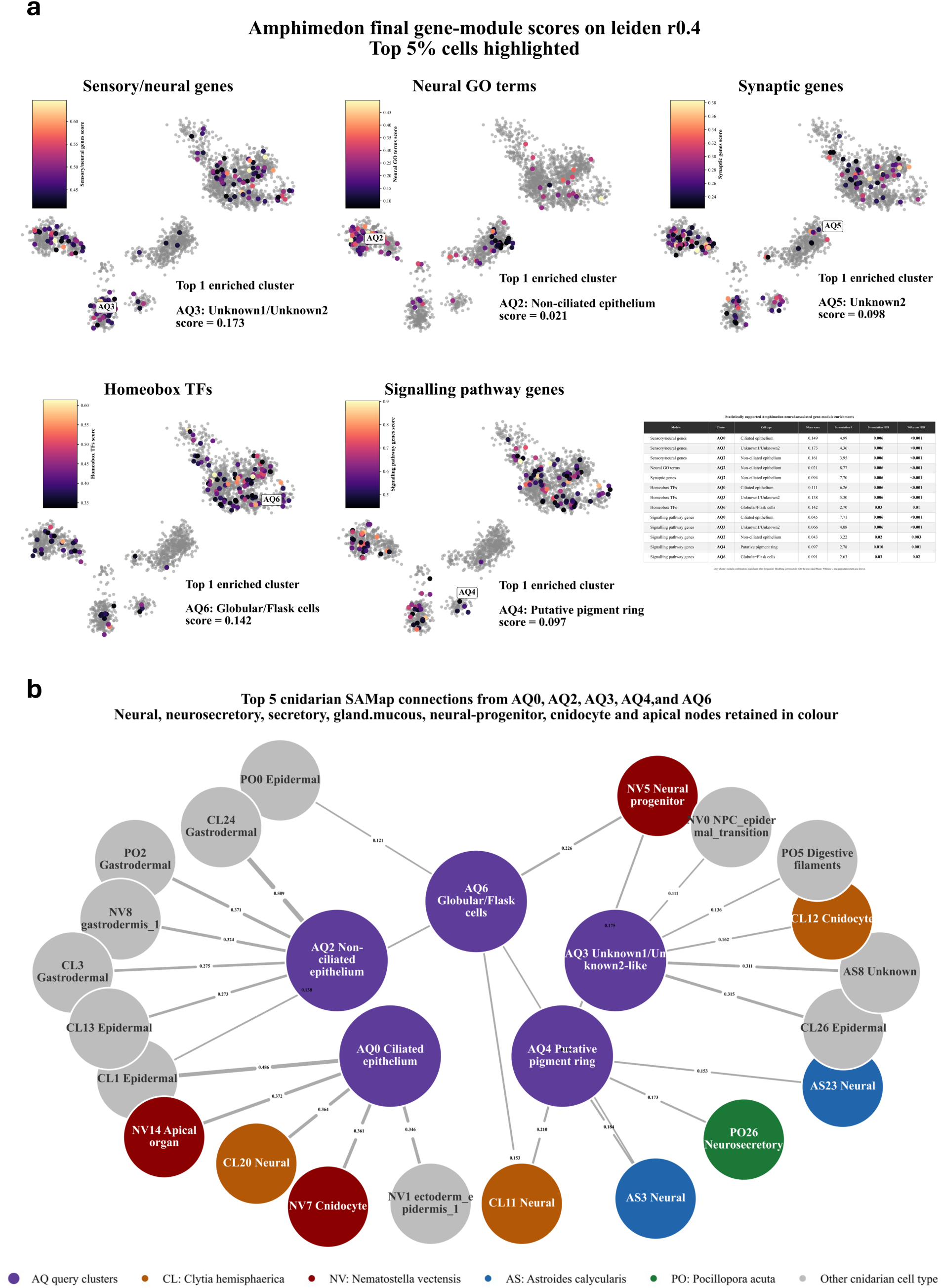
Statistical validation of neural-associated gene module enrichment identifies candidate sensory-neural cell populations in *Amphimedon queenslandica*. **(a)** UMAP visualisations showing the distribution of cells enriched for five neural-associated gene modules compiled from previously published sponge studies^49,61,62^. Gene modules comprise sensory/neural genes, neural Gene Ontology (GO) terms, synaptic genes, homeobox transcription factors and signalling pathway genes. Cells within the top 5% of module scores are highlighted, and the highest-scoring cell clusters for each module are indicated. The accompanying table summarises the statistical validation of module enrichment using Shapiro–Wilk normality tests, one-sided Mann–Whitney U tests and permutation-based null models with Benjamini–Hochberg false discovery rate (FDR) correction. **(b)** SAMap network showing the strongest cross-species transcriptional relationships for statistically supported *Amphimedon* cell populations (AQ0, AQ2, AQ3, AQ4 and AQ6). Each *Amphimedon* cluster is connected to its five strongest cnidarian SAMap matches. Coloured cnidarian nodes represent neural, neurosecretory, secretory, gland-mucous, cnidocyte and apical organ cell types, whereas grey nodes indicate all other cnidarian cell types.

To determine whether these statistically supported *A. queenslandica* larval cell populations exhibited transcriptomic relationships with neural and sensory-secretory cell types in cnidarians, we examined their five strongest cross-species SAMap connections (Fig. 6b). Ciliated epithelium (cluster 0) showed transcriptional similarity to *C. hemisphaerica* neural cluster 20 (0.36) and the *N. vectensis* apical organ cluster 14 (0.37) and cnidocyte cluster 7 (0.36), alongside stronger connections to non-neural epithelial populations. Putative pigment ring cells (cluster 4) displayed broad similarities to cnidarian neural and neural-associated populations, including neural populations in *C. hemisphaerica* (0.21) and *A. calycularis* (0.18 and 0.15), *N. vectensis* neurogenic progenitors (0.17), and a source-annotated neurosecretory population in *P. acuta* (0.17). Globular/flask cells (cluster 6) similarly showed transcriptional similarity to *N. vectensis* neurogenic progenitors (0.23) and *C. hemisphaerica* neural cells (0.15), although several of their strongest connections were to non-neural populations. In contrast, non-ciliated epithelium (cluster 2) and Unknown1/Unknown2 (cluster 3), despite significant enrichment for multiple neural-, sensory-, synaptic-, developmental-or signalling-associated gene modules, did not connect to targeted cnidarian neural or sensory-secretory populations among their five strongest SAMap matches. Collectively, ciliated epithelium (cluster 0), putative pigment ring cells (cluster 4) and globular/flask cells (cluster 6) exhibited both statistically supported gene-module enrichment and cross-species transcriptional similarities to cnidarian neural and sensory-associated cell types, whereas non-ciliated epithelium (cluster 2) and Unknown1/Unknown2 (cluster 3) showed significant gene-module enrichment without corresponding transcriptional similarities among their strongest SAMap connections. These findings suggest that transcriptional programmes associated with neural cell types may have evolutionary antecedents in multiple sponge larval cell populations, including specialised sensory-like and epithelial cells.

### WGCNA reveals cell type-specific transcriptional programmes across cnidarians and sponges

Across the four cnidarian atlases, WGCNA identified between 28 and 40 transcriptional co-expression modules, the majority of which (>75%) exhibited enrichment in only one or two annotated cell populations (Fig. 7a–d). The genes assigned to each co-expression module are provided as Supplementary Data. Co-expression programmes consistently delineated the major larval cell types, including epidermal, gastrodermal, gland/mucous, secretory, cnidocyte, apical, neural and neurosecretory populations. Neural and neurosecretory cell types were characterised by multiple lineage-restricted co-expression programmes, indicating extensive transcriptional diversification within the cnidarian nervous system. Additional modules defined specialised cell populations, including neurogenic progenitors and cnidocyte lineages in *N. vectensis* and proliferative i-cell populations in *C. hemisphaerica*, highlighting the ability of WGCNA to resolve transcriptionally distinct developmental and differentiated cell states across cnidarian larvae.

**Figure 7.**
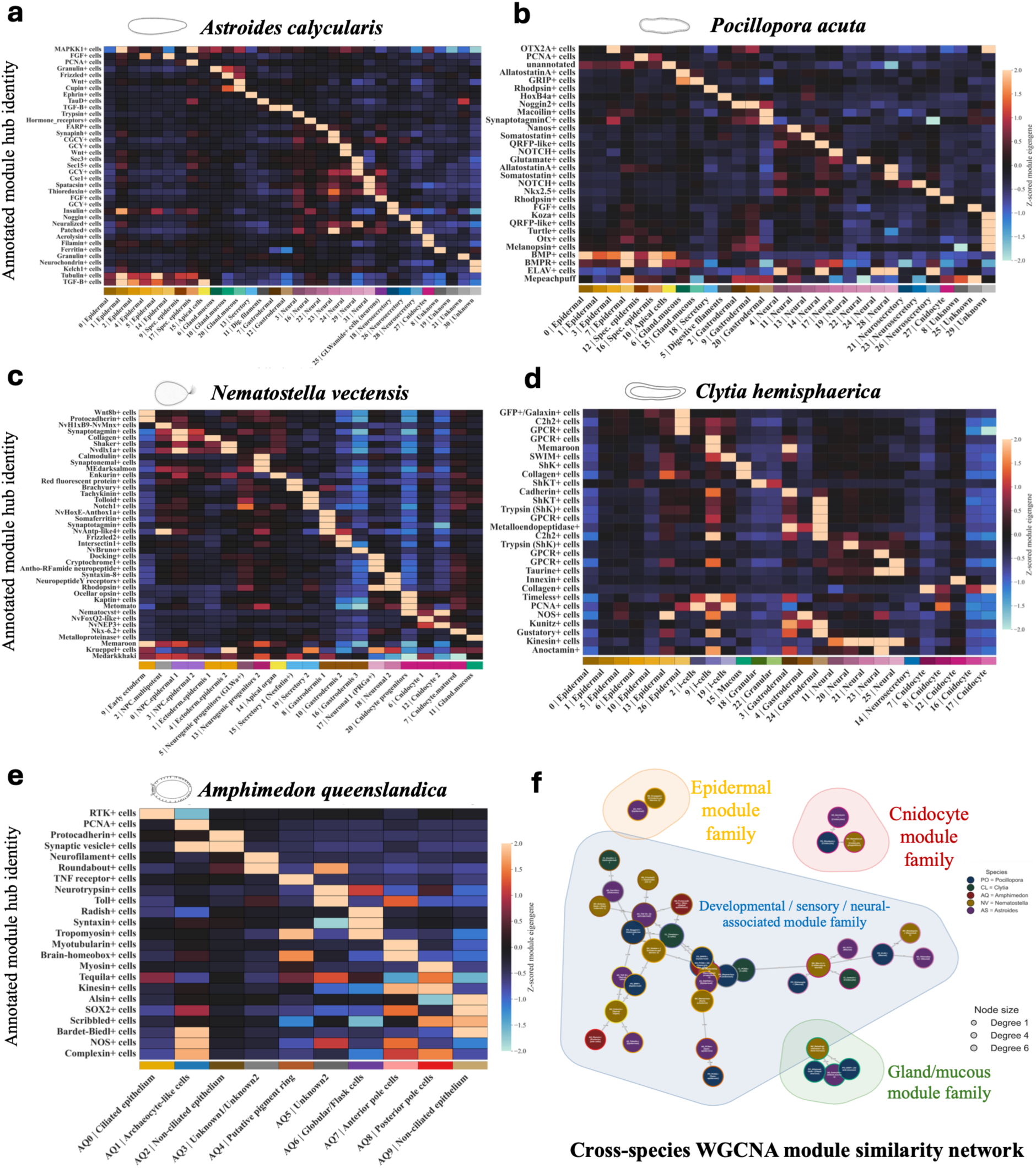
Cell type-specific transcriptional programmes identified by weighted gene co-expression network analysis (WGCNA). (a–e) Heatmaps showing the relative eigengene activity of annotated WGCNA modules across transcriptionally defined cell clusters for **(a)** *A. calycularis*, **(b)** *P. acuta*, **(c)** *N. vectensis*, **(d)** *C. hemisphaerica* and **(e)** *A. queenslandica*. WGCNA modules are annotated according to their highest intramodular connectivity (hub) gene. Columns represent annotated cell clusters, while rows represent WGCNA modules. Colours indicate row-scaled module eigengene expression (Z-score), highlighting cell type-specific enrichment of individual co-expression programmes. The coloured strip beneath each heatmap indicates the broad cell-type identity of each cluster, with related clusters shown as graded shades of a common colour. **(f)** Cross-species WGCNA module similarity network constructed from orthogroup overlap between WGCNA modules. Nodes represent individual modules, labelled according to their annotated hub gene. Node fill colours indicate species of origin, whereas node border colours denote the cell type in which each module exhibits its highest eigengene activity. Edges represent orthogroup-based similarity between modules, and only the 50 strongest pairwise module overlaps are shown.

The *A. queenslandica* atlas likewise exhibited pronounced transcriptional modularity, comprising 23 co-expression modules, 21 of which (91.3%) were enriched in only one or two annotated cell populations (Fig. 7e). Distinct co-expression programmes characterised ciliated and non-ciliated epithelia, archaeocyte-like cells, globular/flask cells, anterior and posterior pole cells, and the putative pigment ring. Unknown-labelled clusters were similarly associated with discrete co-expression programmes, supporting their interpretation as coherent transcriptional cell states rather than unresolved intermediates.

Given the transcriptional diversity of cnidarian neural populations, we further examined the distribution of co-expression programmes across neural and neural-associated cell types. Across all four cnidarians, individual neural populations were characterised by multiple highly restricted modules, alongside broader programmes shared among related neural-associated populations. In *A. calycularis*, most neural and neurosecretory modules were restricted to individual populations, although selected modules were shared between neural and neurosecretory clusters. Similarly, *P. acuta* contained numerous neural population-specific modules, while an *ELAV*-annotated module was enriched across four neural populations and one neurosecretory population. In *N. vectensis*, distinct modules characterised the two differentiated neuronal populations, alongside modules shared between neuronal populations and between neurogenic progenitor and secretory states. *C. hemisphaerica* likewise exhibited both population-specific neural modules and a broader *Kinesin*-annotated module enriched across all five neural populations. Together, these patterns reveal extensive transcriptional diversification among cnidarian neural cell types while also identifying co-expression programmes shared across subsets of neural, neurosecretory and developmental cell states.

Given the central role of larval sensory populations in our comparative framework, we further examined co-expression programmes associated with apical and sensory-like cell populations across the datasets. In the cnidarians, apical cell populations were characterised by distinct developmental and signalling-associated modules. The *A. calycularis* apical population was associated with *TGF-β-*, *FGF-* and *MAPKK1*-annotated modules, while the *P. acuta* apical population was associated with *BMP-* and *BMPR*-annotated modules. In *N. vectensis*, the apical organ population was associated with *Notch1-*, *Enkurin-* and *Synaptotagmin*-annotated modules. Sensory-associated populations in *A. queenslandica* similarly exhibited distinct co-expression programmes: ciliated epithelial cells were associated with receptor tyrosine kinase- and *Tequila*-annotated modules, anterior pole cells with *Brain-homeobox*-, *Myotubularin-*, *nitric oxide synthase-* and *Complexin*-annotated modules, and putative pigment ring cells with *Brain-homeobox-* and *Tropomyosin*-annotated modules. Together, these patterns highlight distinct combinations of co-expression programmes associated with larval apical and sensory-like populations across the cnidarian and sponge datasets.

To investigate whether independently identified co-expression programmes exhibited broader evolutionary relationships, we compared WGCNA modules based on shared orthogroup composition and visualised the 50 strongest pairwise overlaps as a cross-species module similarity network (Fig. 7f). Restricting the analysis to the highest-confidence relationships produced a network comprising 39 modules connected by 50 cross-species edges and representing all five species. Rather than forming a single interconnected network, these modules are resolved into four distinct module families. The largest component contained developmental, sensory, and neural-associated programmes from all species, whereas three smaller components corresponded to conserved gland/mucous, cnidocyte and epidermal-associated co-expression programmes. Proliferative modules formed strong cross-species relationships linking the sponge and cnidarian datasets, while neural-associated programmes repeatedly connected among cnidarian species. Together, these findings identify conserved developmental, sensory-secretory and neural-associated transcriptional programmes that recur across early-diverging metazoans despite the largely species-specific composition of individual WGCNA modules.

## Discussion

Here, we used comparative single-cell transcriptomics to investigate cell-type relationships between the demosponge *Amphimedon queenslandica* and four cnidarian species at the larval stage, with particular focus on sensory-secretory and neural-associated cell types. By integrating cross-species mapping with analyses of transcriptional relationships among cell populations and gene-expression programmes, we examined how these cell types and their underlying molecular programmes have diversified across animal lineages separated by more than 600 million years of evolution^63–65^.

### Conserved cell-type identities underlie extensive transcriptional diversification across cnidarians

Across the four cnidarian species, cross-species mapping recovered broad conservation among corresponding cell-type populations, with neural, secretory, cnidocyte, gland/mucous, epithelial, gastrodermal and apical cell populations repeatedly showing transcriptional similarity to equivalent or closely related cell types across species. These patterns are consistent with the conservation and homology of major cell-type identities across cnidarian lineages despite their substantial evolutionary divergence^52,66–70^. At the same time, neural and secretory populations exhibited considerable transcriptional diversification, retaining broad cross-species similarities while displaying population- and lineage-restricted co-expression programmes, consistent with previous single-cell and developmental studies of cnidarian neural diversity^41,45,46,50^. Apical populations similarly retained a conserved association with epithelial cell types while exhibiting distinct developmental and signalling-associated programmes across species, consistent with the close developmental and transcriptional association of apical sensory populations with larval epithelia described in cnidarians^28,46,71,72^. Beyond these differentiated cell populations, marker-based analyses using the *N. vectensis* neurogenic progenitor signature identified candidate NPC-like populations across the other cnidarian species and in *A. queenslandica*. The transcriptional similarities between the *A. queenslandica* archaeocyte-like population and cnidarian progenitor-like populations may reflect the shared deployment of conserved progenitor-associated molecular programmes across these deeply diverged lineages ^73–75^.

Within-species transcriptional organisation provided further insight into the organisation of neural and secretory cell populations across the cnidarian datasets. Neural and secretory populations clustered closely in the dendrogram analyses and showed substantial connectivity in PAGA, including connections between neural and gland/mucous populations in *A. calycularis* (AS22 – AS10; PAGA connectivity – 1.00) and *P. acuta* (PO19 – PO15; 0.49), and between neural and source-annotated neurosecretory populations in *C. hemisphaerica* (CL21 – CL14; 0.28). Such patterns have a developmental basis in cnidarians, where neural and secretory cell types can arise from shared or closely related progenitor populations regulated by common neurogenic transcriptional programmes ^41,50^. This developmental association was particularly apparent in *N. vectensis*, where neuronal populations showed strong connectivity with neurogenic progenitors, most notably between neuronal cluster 17 and neurogenic progenitor cluster 5 (PAGA connectivity = 0.57), consistent with studies implicating regulators such as *SoxC* and *Insm1* in the differentiation of neural and secretory lineages^45^.

Within cnidarians, taurine-associated programmes provide an additional example of related molecular programmes being deployed in different cellular contexts. A taurine-associated module was linked to neural cells in *C. hemisphaerica*, whereas taurine-associated features occurred in gland/mucous and specialised epithelial populations in *A. calycularis*. Consistent with these patterns, taurine-associated transport and metabolic machinery is concentrated in aboral sensory-neural cells of *C. hemisphaerica* but in specialised aboral epidermal cells of coral larvae, with taurine itself experimentally shown to inhibit larval settlement^46^. Earlier studies in *Hydractinia echinata* also investigated a potential role for taurine in metamorphosis, although their findings were conflicting: an initial study proposed that endogenous taurine may stabilise the larval state, whereas subsequent experiments failed to reproduce the reported decrease in taurine during metamorphosis and instead supported a prominent role for taurine in cellular osmoregulation^76,77^. Given these contrasting findings, we conservatively interpret our results as evidence for the deployment of taurine-associated molecular programmes across distinct cnidarian larval cell types, rather than assigning a conserved physiological function to these programmes.

### Sponge sensory-associated populations share molecular features with cnidarian neural-associated cell types

The conserved yet transcriptionally diverse cnidarian larval cell types provide an important reference for interpreting the position of *A. queenslandica* cell types in the comparative analyses. Several sponge sensory-associated populations exhibited transcriptional similarity to cell types belonging to these conserved cnidarian neural, secretory and sensory-associated groups, suggesting shared components of their molecular programmes. Whereas corresponding cell types across the sampled cnidarians can be considered homologues, the transcriptional similarities between sponge and cnidarian populations instead indicate conservation or shared deployment of molecular programmes across these deeply diverged lineages.

The hypothesis that neurons evolved through the progressive modification of ancestral sensory-secretory cell types has received substantial support from developmental and comparative studies across early-diverging metazoans^78–82^. Our analyses extend this perspective to sponge–cnidarian comparisons, with several *A. queenslandica* sensory-like populations exhibiting transcriptional similarities to cnidarian neural-associated cell types. Cilliated epithelial cells showed SAMap similarities to the *N. vectensis* apical organ (0.37) and neuronal populations (0.31), as well as *C. hemisphaerica* neural cells (0.36). Putative pigment-ring cells similarly mapped to neural populations in *C. hemisphaerica* (0.21) and *A. calycularis* (0.18), neurogenic progenitors in *N. vectensis* (0.17), and a source-annotated neurosecretory population in *P. acuta* (0.17). Globular/flask cells showed their strongest relevant cnidarian similarity to *N. vectensis* neurogenic progenitors (0.23), together with similarity to *C. hemisphaerica* neural cells (0.15).

The relevance of these transcriptional similarities is further supported by previous morphological, functional and molecular studies of sponge larval sensory cell types. In *A. queenslandica*, anterior flask cells represent one of the best-characterised examples, exhibiting ultrastructural features associated with both sensory and secretory functions, including an apical cilium, an F-actin-rich protrusion and abundant secretory vesicles^16^. These cells respond to environmental settlement cues through calcium-dependent signalling and participate in the initiation of metamorphosis^16^. Other specialised populations are distributed throughout the larval epithelium, including globular cells and posterior pigment-ring-associated cells, which have also been implicated in larval sensory and behavioural functions^18,62^. More broadly, sponge genomes encode components of molecular systems associated with neural communication in eumetazoans, including conserved postsynaptic scaffold and signalling components^49,61,83^. Together, these established morphological, functional and molecular characteristics provide important context for interpreting the transcriptional similarities identified here between *A. queenslandica* sensory-associated populations and cnidarian neural-associated cell types.

### Neural-associated molecular programmes are distributed across multiple sponge larval cell types

A notable outcome of our comparative analyses is that neural-associated transcriptional programmes were not restricted to a single sponge sensory-like cell population. Importantly, enrichment of these programmes did not necessarily correspond to broader transcriptional similarity with cnidarian neural populations, as illustrated by AQ2 and AQ3, which showed significant enrichment across several gene modules but lacked corresponding neural-associated similarities among their strongest SAMap connections. In a subset of populations, however, gene-module enrichment coincided with transcriptional similarities to cnidarian neural-associated and secretory populations. Independent neural gene-module analyses therefore indicate that neural-associated programmes are distributed across multiple *A. queenslandica* larval cell populations, while only a subset also exhibit broader transcriptional similarities to cnidarian neural-associated cell types. A similar distributed pattern has been reported in juvenile freshwater sponges, where multiple cell-type families deploy conserved neural gene sets^84^.

Examination of individual genes comprising the sensory/neural gene module (Fig. 6a; Supplementary Fig. S6) provided further insight into the evolutionary history of molecular components associated with nervous-system function. Several of these genes belong to evolutionarily conserved gene families present across sponges, cnidarians and bilaterians, although their cellular deployment and functions may have diverged across these lineages^19,84,85^. *Cry2* was most strongly expressed in putative pigment-ring cells, consistent with its proposed role as a blue-light-sensitive candidate photoreceptor in the *A. queenslandica* larval photoreceptive system^85^, while cryptochromes are also conserved in cnidarians and bilaterians, where they function variously as direct environmental light sensors^86,87^, or as core transcriptional repressors in the endogenous circadian clock^88,89^.

The Atonal-related transcription factor *bHLH1* was concentrated in globular/flask cells, while related bHLH factors regulate neural and sensory cell differentiation in cnidarians and bilaterians^19,90,91^. Similarly, *Homer* and GRIP were most strongly expressed in globular/flask and putative pigment-ring cells, respectively (Supplementary Fig. S7); homologues of these scaffolding components are highly conserved across cnidarians and bilaterians. However, while *Homer* has a demonstrated postsynaptic localisation in cnidarians, the direct functional integration of *GRIP* into postsynaptic receptor scaffolds is primarily characterised in bilaterians^36,92,93^. By contrast, *ELAV*, an evolutionarily conserved RNA-binding protein that served as a classic marker of postmitotic, differentiated neuronal identity in both cnidarians and bilaterians^44,94,95^, was only sparsely expressed and was not concentrated in these sensory-associated sponge populations (Supplementary Fig. S7). Together, these patterns suggest that molecular components subsequently incorporated into sensory and neural systems were already present early in animal evolution, while their differing deployment in *A. queenslandica* indicates that the cellular contexts in which these ancient components function have undergone substantial evolutionary diversification^32,61,84^.

At the within-species level, transcriptional organisation in *A. queenslandica* placed the sensory-associated populations within a broader epithelial and sensory group. Globular/flask cells showed PAGA connectivity with ciliated and non-ciliated epithelial populations (0.19 and 0.17, respectively), while putative pigment-ring cells mostly strongly connected with posterior pole cells (0.24). This organisation is consistent with the distributed sensory architecture of sponge larvae, in which specialised ciliated and pigment-associated cells are integrated within the larval epithelium and mediate environmental sensing and behaviour in the absence of a canonical nervous system²¹. Thus, although these sponge populations share transcriptional similarities with cnidarian neural-associated cell types in our cross-species analyses, their within-species organisation remains closely integrated with the epithelial and sensory architecture of the sponge larva. Future studies combining detailed ultrastructural analyses, such as electron microscopy reconstructions, with investigations of intercellular signalling, including ligand-receptor interactions and potential volume transmission, could help clarify how these specialized sponge cell populations communicate and coordinate sensory responses.

The weighted gene co-expression network analysis (WGCNA) identifies groups of genes with correlated expression patterns, allowing conserved or cell-type-associated transcriptional programmes to be examined beyond individual genes^96,97^. The WGCNA analyses further revealed that this distributed pattern extends to specific components of sensory and secretory signalling machinery. An NO/NOS-associated module was linked to several *A. queenslandica* populations, including anterior pole cells, while previous experimental work has localised *NOS* expression and NO production to larval globular cells and demonstrated that NO signalling promotes settlement and metamorphosis in *A. queenslandica*^18^. In *C. hemisphaerica*, *NOS* is enriched in QYFamide sensory-neural cells at the aboral pole, which also express neural and secretory-associated genes including *ELAV* and synaptotagmin^46^, indicating that NO signalling is deployed within specialised larval sensory-associated cellular contexts in both sponges and cnidarians. Components of vesicle-release machinery showed a similarly distributed organisation in *A. queenslandica*, with *Complexin* associated with anterior and posterior pole cells, *Syntaxin* with globular/flask cells, and an SV2-related protein with archaeocyte-like and non-ciliated epithelial populations. Related secretory machinery, including *Syntaxin*, *Synaptotagmin* and *UNC-13*-associated components, is also deployed in specialised cell populations of other sponges^84^, while our cnidarian datasets recovered multiple synaptic and vesicle-associated components within neural-associated populations, including *Synaptotagmins*, *Synapsin* and *Syntaxin* in *N. vectensis* and *Synapsin*/*SV2*-related features in *P. acuta*. Together, these patterns support the distributed deployment of sensory- and secretory-associated molecular machinery across sponge larval cell types.

### Sensory-secretory molecular programmes inform scenarios of early neuronal evolution

Taken together, our findings are consistent with an evolutionary scenario in which neural-associated molecular programmes were distributed across multiple ancestral sensory-secretory cell types, rather than being assembled within a single ancestral cell type. Subsequent diversification and reorganisation of these cellular programmes may have contributed to the emergence of the diverse neural and other sensory-secretory cell types found in modern animals. This interpretation is compatible with both stepwise models of neuronal evolution and scenarios involving multiple evolutionary origins of neuronal cell types through the recruitment of pre-existing sensory, secretory and signalling machinery^81,98^.

Despite the power of comparative single-cell transcriptomics for investigating cell-type evolution, several limitations should be considered. Transcriptional similarity alone cannot establish direct cell-type homology, as shared molecular programmes may reflect common ancestry, repeated co-option of ancestral regulatory modules or convergent evolution^99–101^. Furthermore, because SAMap identifies cross-species transcriptional similarities through homologues genes, these comparisons may not fully capture lineage-specific transcriptional programmes. Consequently, the integration of sponge larval cell populations within cnidarian communities may reflect shared molecular programmes while overlooking additional transcriptional features that distinguish sponge cell-type identities. Evolutionary interpretation also remains dependent on taxon sampling; incorporating additional early-diverging animal lineages at informative phylogenetic positions could help resolve when sensory, secretory and neural-associated programmes emerged. Comparisons of independently generated single-cell datasets are further affected by differences in experimental design, developmental staging, tissue dissociation, sequencing and computational processing. Greater adoption of shared reporting standards and detailed experimental and analytical metadata would facilitate comparisons across studies, species and laboratories^102^. Further subclustering of the *A. queenslandica* larval dataset could also determine whether the neural-associated programmes distributed across multiple clusters are concentrated within smaller, transcriptionally distinct cell populations that are not resolved at the current clustering resolution. Finally, the transcriptional similarities identified here provide specific hypotheses that can be tested through spatial, developmental and functional approaches to determine whether shared molecular programmes contribute to comparable cellular functions across species.

In conclusion, our comparative analyses provide new insight into the evolutionary assembly of sensory-secretory and neural-associated molecular programmes across early-diverging metazoans. By integrating complementary comparative transcriptomic approaches across free-swimming sponge and cnidarian larvae, we identified transcriptional similarities between multiple *A. queenslandica* sensory-like cell populations and cnidarian neural-associated and secretory cell types, alongside shared sensory, secretory, signalling and developmental molecular programmes. Neural-associated features were distributed across several sponge sensory-like populations, supporting the possibility that the diversity of modern neural and sensory-secretory cell types emerged through the diversification and reorganisation of molecular programmes deployed across multiple ancestral sensory-secretory cell types. Together, these findings provide a comparative perspective on how the molecular machinery underlying sensory integration and intercellular communication may have contributed to the emergence and diversification of specialised neuronal cell types and nervous systems.

## Acknowledgements

We thank the researchers who generated and made publicly available the single-cell transcriptomic datasets used in this study. We also thank Melisa Chan for her assistance with manuscript formatting.

## Author contributions

Conceptualisation: V.M.; Methodology: I.L., J.S.; Software: I.L.; Validation: J.S.; Formal analysis: I.L.; Investigation: I.L.; Resources: I.L., E.W., V.M., J.S.; Data curation: I.L.; Writing – original draft: I.L.; Writing – review & editing: I.L., E.W., V.M., J.S.; Visualisation: I.L.; Supervision: E.W., V.M., J.S.; Project administration: E.W., V.M.; Funding acquisition: E.W., V.M.

