## Supplementary figures and images for "Comparative larval single-cell transcriptomics provides insights into the evolutionary origins of sensory-neural programmes in Porifera and Cnidaria"

### Supplementary Data

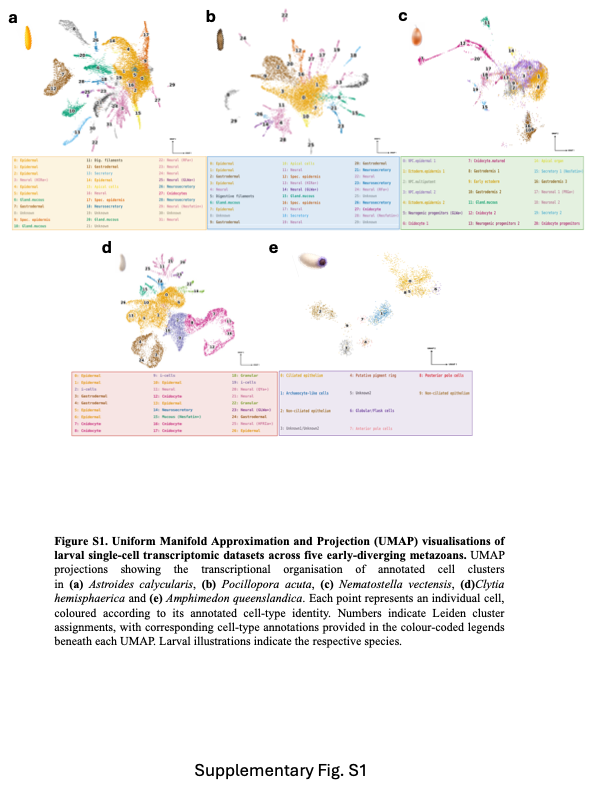

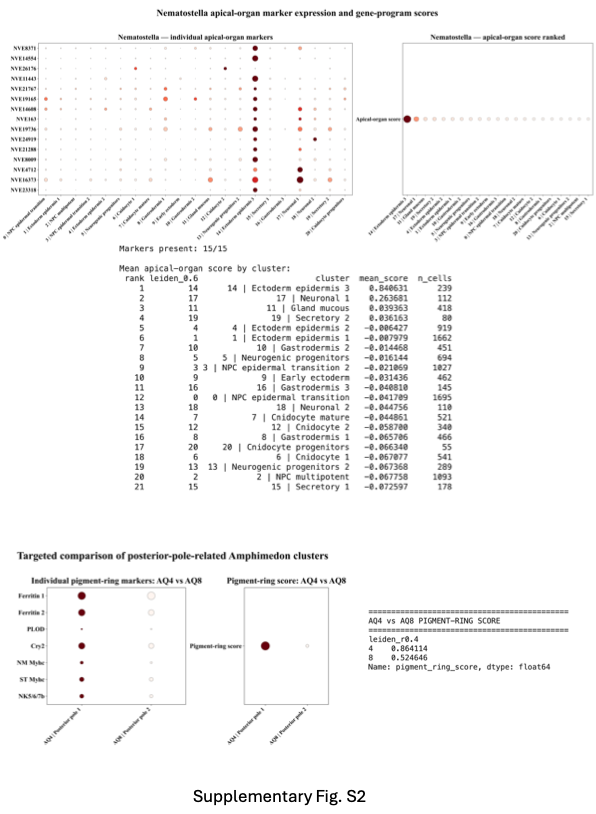

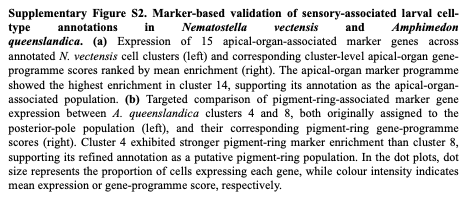

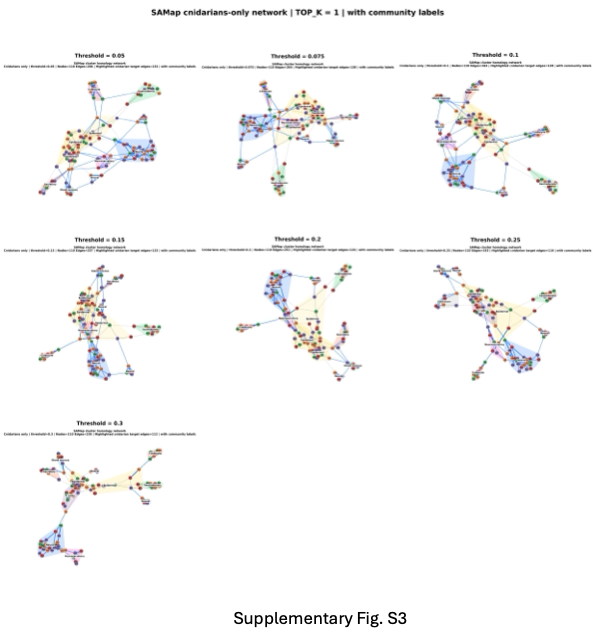

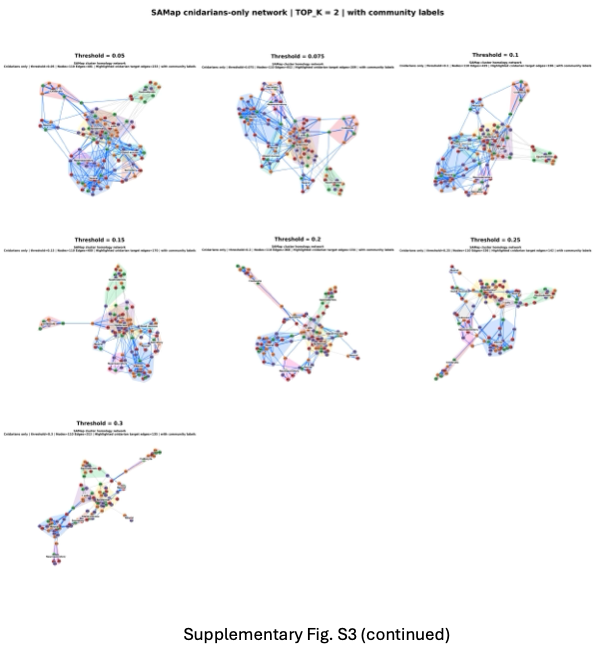

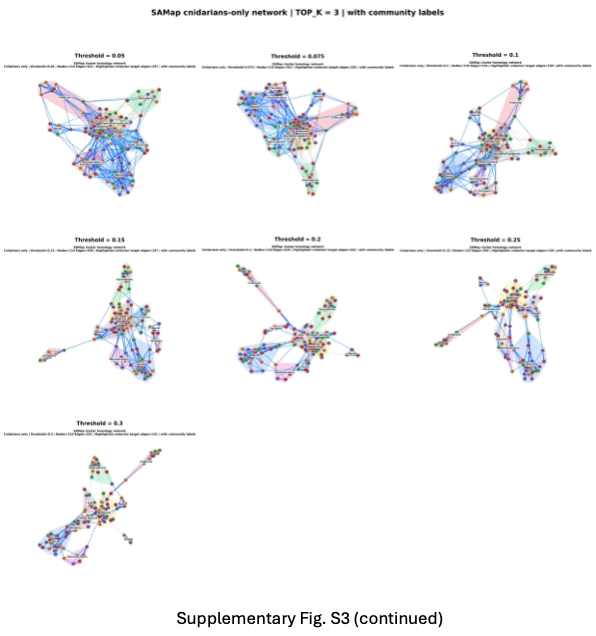

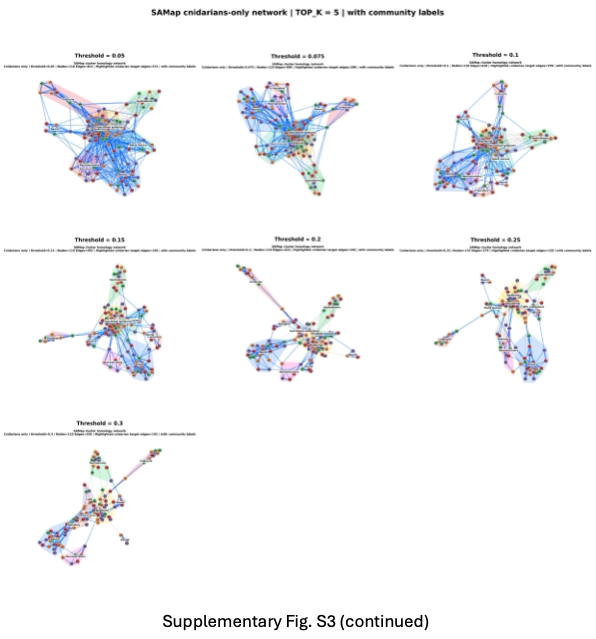

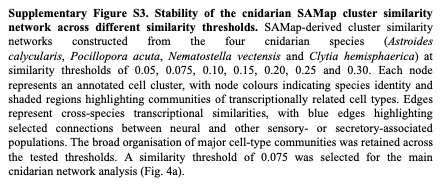

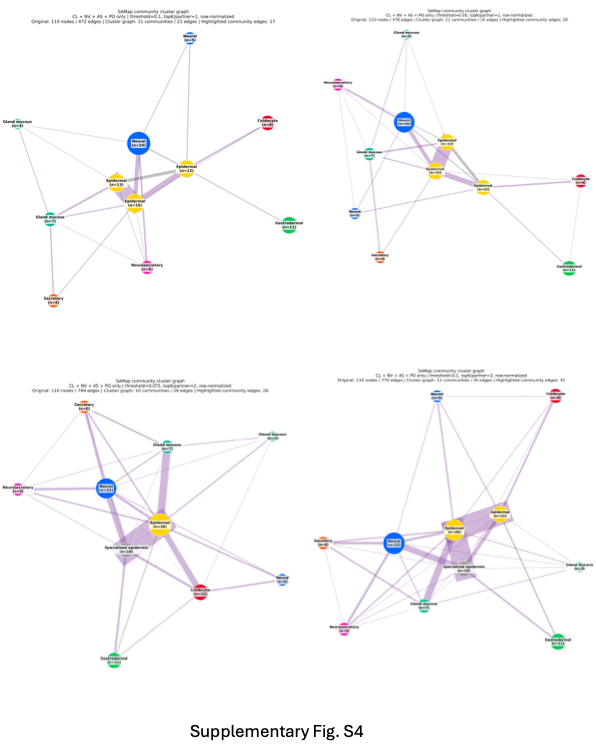

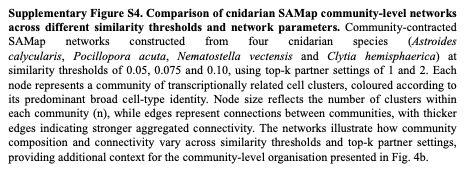

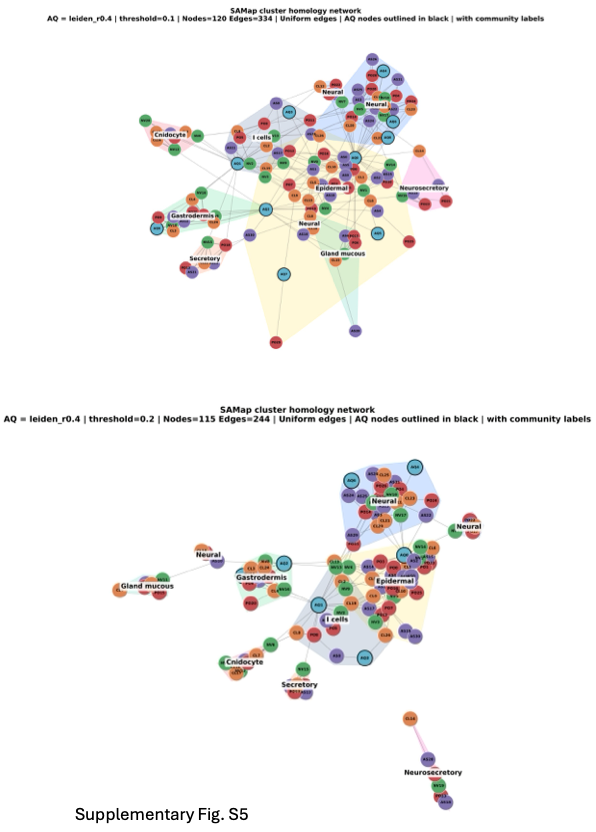

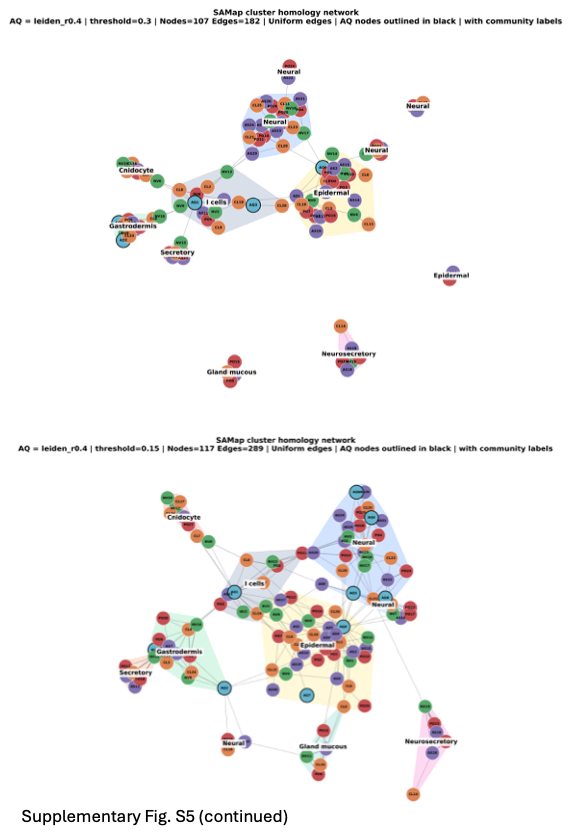

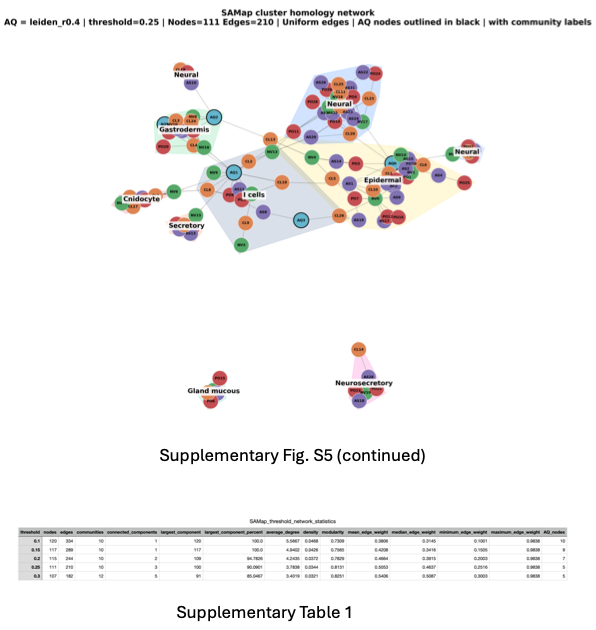

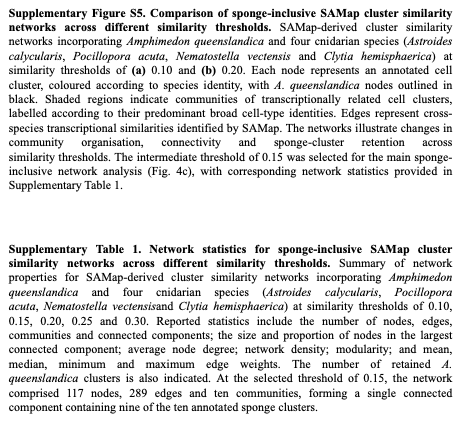

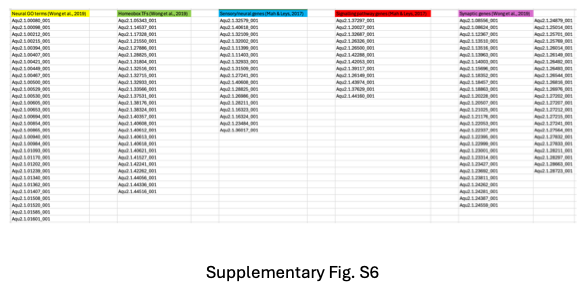

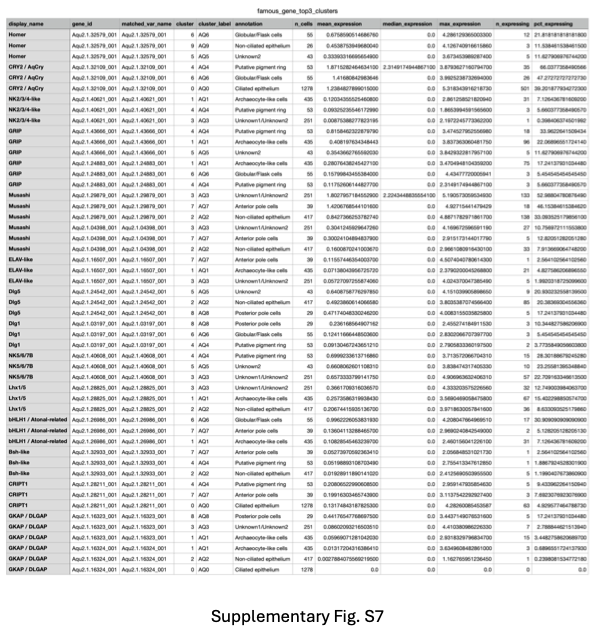
